# Longitudinal functional connectivity during rest and task is differentially related to Alzheimer’s pathology and episodic memory in older adults

**DOI:** 10.1101/2025.05.16.654628

**Authors:** Larissa Fischer, Jenna N. Adams, Eóin N. Molloy, Jennifer Tremblay-Mercier, Jordana Remz, Alexa Pichet Binette, M. Natasha Rajah, Sylvia Villeneuve, Anne Maass, PREVENT-AD Research Group

**Affiliations:** German Center for Neurodegenerative Diseases (DZNE), Magdeburg, Germany; Department of Neurobiology and Behavior, University of California, Irvine, USA; Department of Radiology & Nuclear Medicine, Faculty of Medicine, Otto von Guericke University, Magdeburg, Germany; StoP-AD Centre, Douglas Mental Health Institute Research Centre, Montréal, Canada; Department of Physiology and Pharmacology, Université de Montréal, Montréal, Canada; Centre de Recherche de l’Institut Universitaire de Gériatrie de Montréal, Montréal, Canada; Clinical Memory Research Unit, Department of Clinical Sciences Malmö, Lund University, Lund, Sweden; Department of Psychiatry, McGill University, Montréal, Canada; Department of Psychology, Toronto Metropolitan University, Toronto, Canada; Institute for Biology, Otto von Guericke University, Magdeburg, Germany

**Keywords:** Aging, Alzheimer’s disease, fMRI, Functional Connectivity, Episodic Memory, *APOE4*

## Abstract

Changes in functional connectivity (FC) strength involving the medial temporal lobe (MTL) and posteromedial cortex (PMC) are related to early Alzheimer’s pathology and alterations in episodic memory performance in cognitively unimpaired older adults, but their dynamics remain unclear. We examined how longitudinal changes in FC involving MTL and PMC during resting-state, episodic memory encoding, and retrieval relate to subsequent amyloid- and tau-PET burden, longitudinal episodic memory performance, and the *APOE4* genotype in 152 cognitively unimpaired older adults from the PREVENT-AD cohort. We found *APOE4-* and fMRI paradigm-dependent associations of change in FC strength with pathology burden and change in episodic memory performance. Decreasing FC over time, or “hypoconnectivity”, within PMC during rest in *APOE4* carriers and during retrieval in *APOE4* non-carriers was related to more amyloid and tau, respectively. Conversely, increasing FC over time, or “hyperconnectivity”, within MTL during encoding in *APOE4* carriers and between MTL and PMC during retrieval independent of *APOE4* status was related to more tau. Further, increasing FC between MTL and PMC during rest, unlike during encoding, was beneficial for episodic memory. Our study highlights that pathology-related episodic memory network changes manifest differently during rest and task and have differential implications for episodic memory trajectories.

## Introduction

Episodic memory is one of the first cognitive functions that declines with aging^1^ and Alzheimer’s disease (AD)^2^. Co-activation of medial temporal lobe (MTL) and posteromedial cortex (PMC) regions, which can be measured using functional magnetic resonance imaging (fMRI) and functional connectivity (FC) analysis, plays a critical role in supporting episodic memory encoding and retrieval^3–5^. In cognitively unimpaired older adults, cross-sectional studies reported connectivity alterations in this episodic memory network of MTL and PMC regions during resting-state (task-free) fMRI, particularly lower connectivity with aging^6,7^ and both higher an lower connectivity with early AD pathology^8–10^. However, most findings are based on resting-state fMRI, as only a few studies have examined FC of the episodic memory network in older adults during memory encoding or retrieval, or across multiple fMRI paradigms that combine rest and task states^11–13^. Moreover, it is unclear which network changes are dysfunctional and related to detrimental processes, i.e. production and spread of AD pathology, and which network changes could be beneficial or compensatory for memory performance^14,15^.

Accumulation of AD pathology, consisting of tau tangles starting within the MTL and amyloid-beta (Aβ) plaques occurring early within the PMC^16,17^, begins while older adults are still cognitively unimpaired^18,19^. The vulnerability of the MTL and the PMC to AD pathology on the one hand, and the importance of these regions for episodic memory on the other hand, highlights the need to better understand how changes in the functional dynamics of MTL and PMC relates to AD risk factors and pathology in cognitively unimpaired older adults^20^. AD pathology can be measured via positron emission tomography (PET), and is associated with higher neural activity in cognitively unimpaired older adults^15,21,22^. Further, findings from predominantly resting-state FC (rsFC) studies suggest complex dynamics between connectivity, AD pathology, and episodic memory performance in older individuals. Increasing rsFC strength, or “hyperconnectivity”, within the MTL and between the MTL and the PMC could drive tau spread across the brain and thus represent an important factor in disease progression^23–25^. On the other hand, decreasing rsFC strength, or “hypoconnectivity”, and hypometabolism within the PMC have been linked to the accumulation of Aβ pathology and could represent a disconnection within the PMC^26–28^. Hypoconnectivity within the PMC at rest has been further related to cognitive decline in previous studies^27,29–31^, while hyperconnectivity between the MTL and the PMC at rest has been suggested to be detrimental for cognition^25,32^.

However, other studies also found a positive association^9,33,34^ or no relationship^35^ between MTL-PMC rsFC strength and cognitive performance in older adults.

The *APOE4* genotype is the strongest genetic risk factor for sporadic AD and is correlated with Aβ accumulation^36,37^. Differences or longitudinal changes in rsFC strength involving the MTL and the PMC have also been reported in cognitively unimpaired older *APOE4* carriers relative to non-carriers^38,39^. For instance, Salami and colleagues reported that hlongitudinal plasma p-tau_181_ increases were paralleled by elevated local hippocampal rsFC strength and subsequent reduction of hippocampus encoding-related activity in *APOE4* carriers^39^. Further findings suggest that lower rsFC strength between episodic memory regions in *APOE4* carriers is beneficial with regard to cognition ^30,38^.

Age- and AD-related longitudinal changes in FC strength in cognitively unimpaired older adults have been investigated primarily using rsFC^40,41^. Only a few studies have investigated FC in parallel during rest and during the performance of cognitive tasks, such as mnemonic discrimination and delayed match-to-sample paradigms^11,32^. These cross-sectional studies found differential associations of FC during rest and task states with AD pathology. It is, however, still unclear whether longitudinal network dynamics across rest, encoding, and retrieval show similarities or differences. Assessing network dynamics over time during different fMRI paradigms and their associations with pathology and cognitive performance measures could help to better understand the complex functional brain changes that are involved in aging and disease, which cross-sectional studies cannot offer^42^.

Thus, in the present preregistered study we investigated (a) how longitudinal changes in FC within and between MTL and PMC regions are related to AD pathology and the *APOE4* genotype in the aging brain, (b) how these changes are related to changes in episodic memory performance, and (c) whether similar changes in FC occur during resting-state and episodic memory task fMRI. We assessed FC at rest, during object-location episodic memory encoding and retrieval, subsequent Aβ- and tau-PET, *APOE4* status, and cognitive data from the PREVENT-AD cohort of cognitively unimpaired older adults. We focused on FC strength within the MTL, within the PMC, and between the MTL and the PMC. Further information regarding the project is provided in the preregistration focusing on resting-state^43^ and task^44^ fMRI. “FC” in the following hypotheses refers to both FC during rest and task states.

Regarding longitudinal FC strength of the episodic memory network and AD pathology, we hypothesized that i) FC shows an age-related decrease, however, ii) FC shows an increase related to higher subsequent Aβ and tau burden, especially in *APOE4* carriers. Regarding longitudinal FC strength of the episodic memory network and longitudinal episodic memory performance, we hypothesized that iii) increasing FC strength could be an initial beneficial or compensatory process if predicting maintained episodic memory performance or a detrimental process if predicting decline in performance.

## Results

### Demographics

One hundred fifty-two cognitively unimpaired older adults (63±5years, 102 female, 59 *APOE4*), from the Pre-symptomatic Evaluation of Experimental or Novel Treatments for Alzheimer’s Disease (PREVENT-AD) cohort^45,46^ were included. Demographics are presented in Table 1. In the current study, we focused on a subsample of PREVENT-AD participants with available longitudinal fMRI data and cross-sectional PET data. In particular, participants completed at least a baseline session of resting-state and episodic memory task fMRI, cognitive testing, and up to four years of follow-up sessions. Additionally, participants underwent PET imaging to measure Aβ using ^18^F-NAV4694 (NAV) and tau using ^18^F-flortaucipir (FTP). PET was conducted on average 63 months after the baseline session (see Figure 1a for an overview).

**Figure 1.**
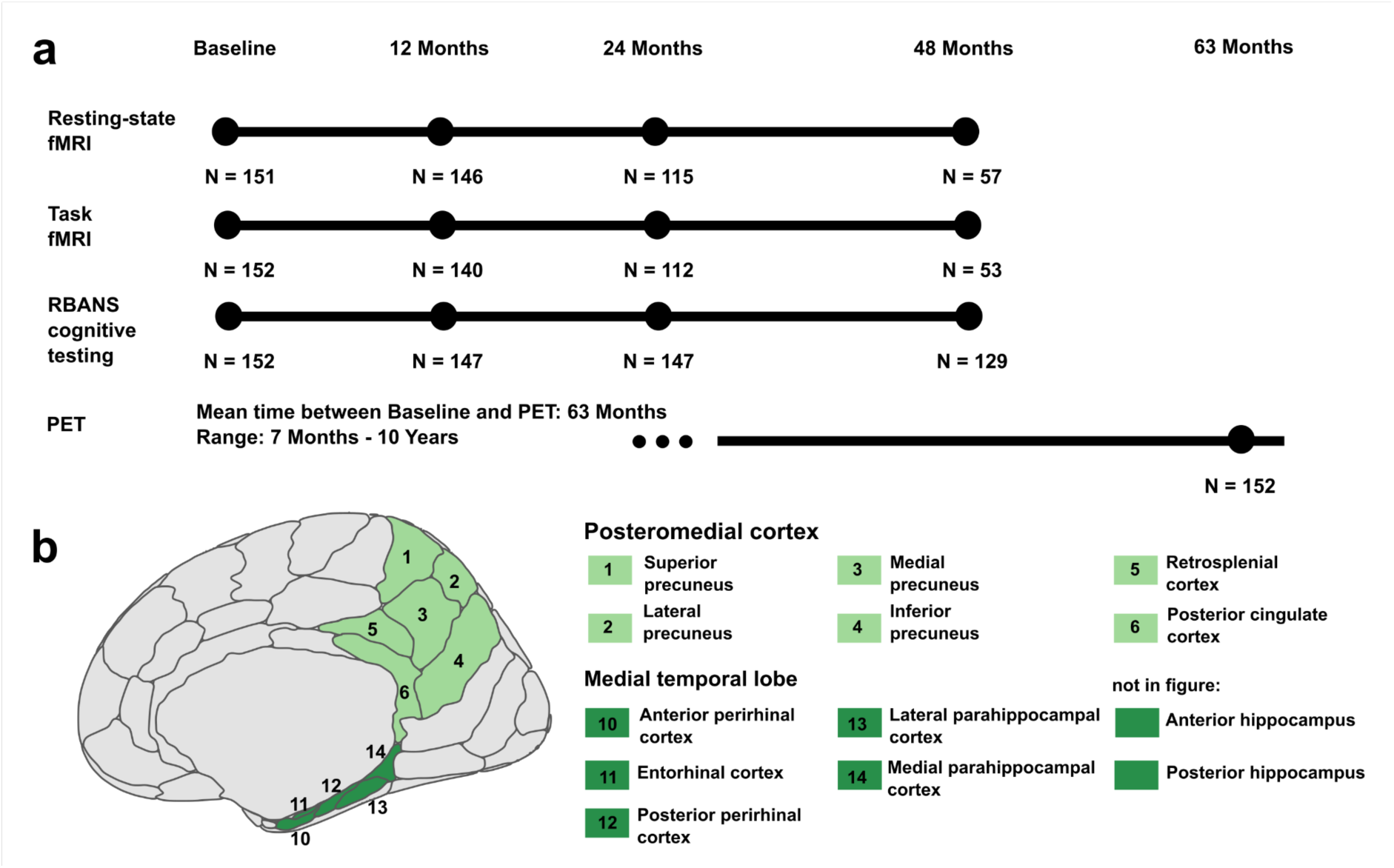
Study design and a priori defined regions of interest (ROIs). **a)** Each participant underwent at least one baseline resting-state and task fMRI session and between 1 and 3 follow-up fMRI examinations, with the last scan taking place 4 years (48 months) after baseline. Similarly, neuropsychological RBANS assessments were conducted at baseline and at follow-up sessions. Participants also underwent PET scanning to quantify amyloid using ^18^F-NAV4694 and tau using ^18^F-flortaucipir. This took place between 7 months and 10 years after the baseline session. Numbers (N) of data for each session and modality describe the cohort after exclusions due to MRI quality control. **b)** The posteromedial cortex (PMC) ROI (light green) consisted of the retrosplenial cortex, posterior cingulate cortex, and four subregions of the precuneus. The medial temporal lobe (MTL) ROI (dark green) consisted of the anterior and posterior perirhinal cortex, entorhinal cortex, lateral and medial parahippocampal cortex, and anterior and posterior hippocampus. Regions were defined using the Brainnetome atlas. fMRI, Functional Magnetic Resonance Imaging; RBANS, Repeatable Battery for the Assessment of Neuropsychological Status; PET, Positron Emission Tomography.

**Table 1:** Final sample demographics

| | *APOE4* non-carrier<br>(N = 93) | *APOE4* carrier<br>(N = 59) | t/ $\chi^2$/ OR | *p* |
| --- | --- | --- | --- | --- |
| Age (years) | 62.7 (4.1) | 63.9 (4.7) | 1.699 | 0.092 |
| Education (years) | 15.5 (3.4) | 15.3 (3.2) | 0.368 | 0.713 |
| Sex (male/ female) | 29/ 64 | 21/ 38 | 0.150 | 0.699 |
| Amyloid PET burden (SUVR) | 1.22 (0.22) | 1.42 (0.36) | -4.366 | **<0.001** |
| Amyloid positive (N) | 7 | 23 | 20.606 | **<0.001** |
| Tau PET burden (SUVR) | 1.04 (0.1) | 1.07 (0.14) | -1.547 | 0.125 |
| Tau positive (N) | 2 | 2 | 1.591 | 0.642 |
| Time baseline to PET (years) | 5.11 (2.3) | 5.49 (2.2) | -0.996 | 0.321 |
| Average follow-up time for MRI<br>(years) | 3.22 (0.81) | 2.98 (0.82) | 1.788 | 0.076 |

Mean (standard deviation) or number of participants are presented. We conducted t-tests, chi-squared tests (for sex and amyloid-positive participants) and a fisher's exact test (for tau-positive participants) for *APOE4* non-carrier versus *APOE4* carrier. Three *APOE4* carriers were homozygous. All participants identified as either female or male. For Aβ PET, global neocortical Aβ was investigated. An amyloid positivity threshold of an SUVR value of 1.39 was provided by the PREVENT-AD Research Group. For tau PET, entorhinal tau was investigated. A tau positivity threshold of an SUVR value of 1.30 was provided by the PREVENT-AD Research Group. *APOE*, apolipoprotein E; N, number; OR, odds ratio; Aβ , amyloid-beta; PET, positron emission tomography; SUVR, standardized uptake value ratio.

### Functional connectivity change over time and relation to *APOE4* status, age, sex, and education

Values for FC strength between each pair of regions of interest (ROIs) were derived by extracting the respective BOLD time-series and conducting ROI-to-ROI functional connectivity analysis. See Figure 1b for details on the 13 included ROIs from the Brainnetome atlas^47^. We calculated the mean FC strength for each participant and session for three meta-ROIs, i.e. (i) all ROIs within MTL, ii) all ROIs within PMC, and iii) between MTL and PMC ROIs. We followed this procedure for the resting-state fMRI session and the encoding and retrieval session from an object-location episodic memory fMRI task^48^. During encoding, participants were presented with 48 objects on either the left or right side of the screen and indicated the side via button press. During retrieval 20 minutes later, participants were presented with 48 old (previously encoded) and 48 new (not previously encoded) objects. They were instructed to give one of the four following forced-choice retrieval responses: familiar object, remembered left, remembered right, or new object.

First, we investigated whether meta-ROI (i.e. within MTL, within PMC and between MTL and PMC) FC strength changed over time for the different fMRI paradigms (resting-state, rsFC; encoding task, encoding-FC; retrieval task, retrieval-FC). We included a time by *APOE4* group interaction, age at baseline, sex, and education as covariates in all models. Linear mixed models revealed that rsFC decreased within MTL (β=-0.10 [95% CI -0.19, -0.01], t=-2.199, *p*=0.029), within PMC (β=-0.13 [95% CI -0.21, -0.04], t=-2.904, *p*=0.004), and between MTL and PMC (β=-0.09 [95% CI -0.18, -0.00], t=-1.988, *p*=0.048) over time. In contrast, encoding- FC and retrieval-FC did not change significantly over time (all *p*>0.05) in the whole sample.

There were no significant effects of *APOE4* group or time by *APOE4* group interactions (all *p*>0.05). Higher baseline age was related to lower retrieval-FC within MTL (β=-0.18 [95% CI -0.32, -0.05], t=-2.728, *p*=0.007). Further, in all models except for two — encoding-FC and retrieval-FC between the MTL and PMC — significant sex effects were observed, with females showing lower FC strength than males. See Supplementary Tables S1-9 for all models.

Exploratively, we compared the trajectories of FC strength between resting-state, encoding and retrieval. The slopes of FC strength over time within MTL were positively correlated between all three fMRI paradigms and changes in FC strength within PMC were positively correlated between resting-state and encoding (r-values between 0.30 and 0.67). Interestingly, changes in MTL-PMC FC strength during rest versus encoding were negatively correlated (r = -0.19). See Table 2 for statistics.

**Table 2:** Correlations of slopes of functional connectivity strength between paradigms

|  | resting state - encoding | resting state - retrieval | encoding - retrieval |
| --- | --- | --- | --- |
| Within MTL | $r = 0.301, t = 3.863, p < 0.001$ | $r = 0.301, t = 3.864, p < 0.001$ | $r = 0.668, t = 11, p < 0.001$ |
| Within PMC | $r = 0.324, t = 4.192, p < 0.001$ | $r = 0.041, t = 0.497, p = 0.631$ | $r = 0.124, t = 1.532, p = 0.191$ |
| Between MTL - PMC | $r = -0.185, t = -2.307, p = 0.040$ | $r = 0.039, t = 0.481, p = 0.631$ | $r = 0.093, t = 1.139, p = 0.330$ |

The correlations of the slopes over time of FC strength between resting-state, encoding and retrieval were assessed. The slopes of FC strength over time correlated between all 3 fMRI paradigms within MTL and for resting state and encoding within PMC. FDR-corrected p-values are reported. FDR correction for multiple comparisons was applied due to the exploratory nature of this analysis. Significant results after correction are highlighted in bold. FC, functional connectivity; FDR, False-discovery-rate; MTL, medial temporal lobe; PMC, posteromedial cortex.

### Relationship between functional connectivity change and subsequent Aβ and tau burden

We then investigated the association of the slopes of MTL and PMC FC strength and subsequent AD pathology (i.e. Aβ- and tau-PET burden) using linear models including the interaction of FC slope by *APOE4* group, age, sex, education, and time between baseline session and PET.

Regarding global neocortical Aβ burden, we found an effect of rsFC slope within PMC that differed by *APOE4* group (β=-0.42 [95% CI -0.81, -0.02], t=-2.100, *p*=0.038; see Figure 2a and Supplementary Table S10). Specifically, *APOE4* carriers showed a steeper decrease of rsFC within PMC with more Aβ (β=-0.35 [95% CI -0.70, -0.01], t=-1.970, *p*=0.054; see Supplementary Table S11), while *APOE4* non-carriers had no association between rsFC slope and Aβ (β=0.01 [95% CI -0.27, 0.29], t=0.071, *p*=0.944; see Supplementary Table S12).

**Figure 2.**
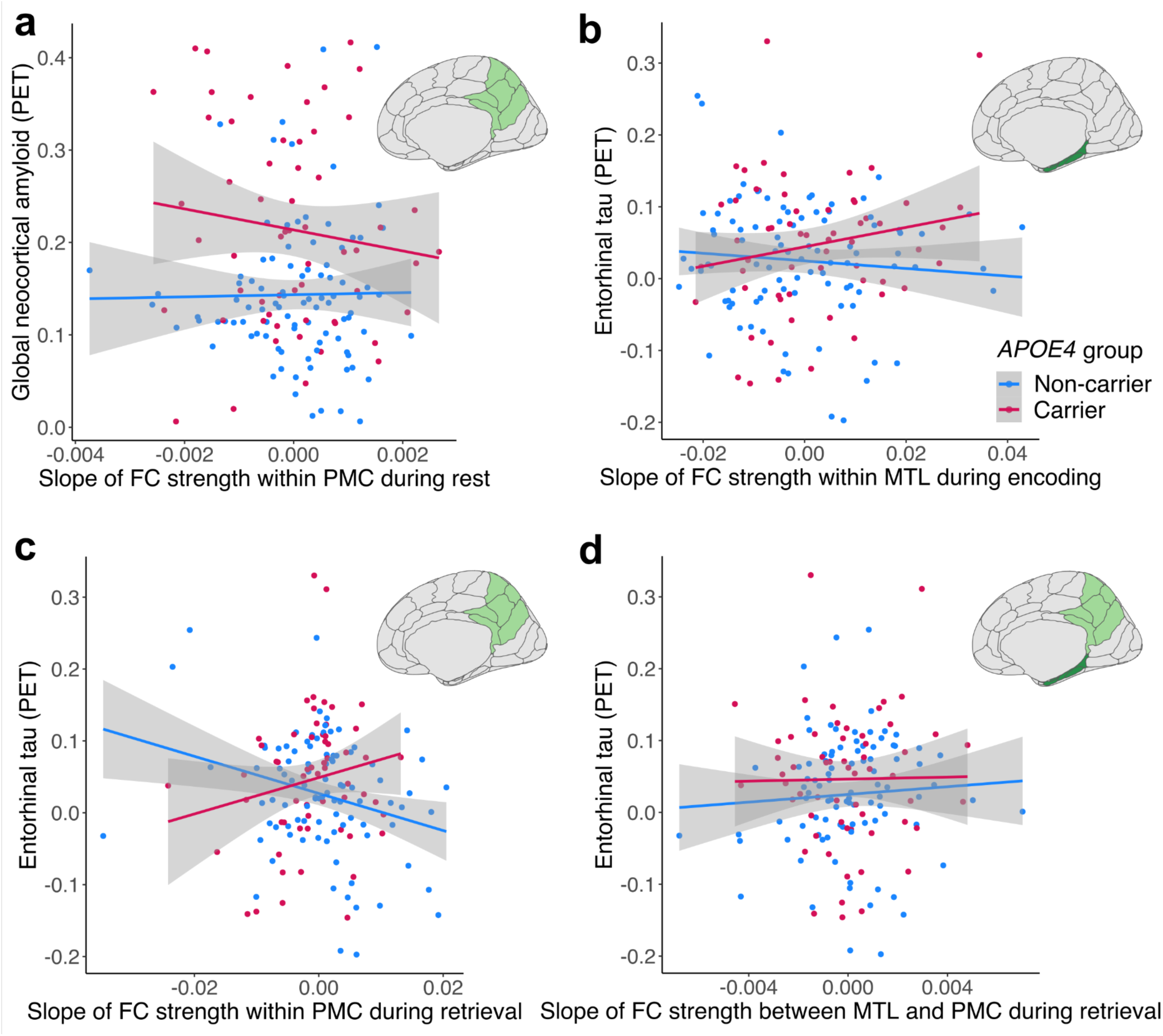
Change in functional connectivity (FC) strength during rest, encoding, and retrieval over time and the relationship with Alzheimer’s pathology measured via PET imaging. **a)** The slope over time of mean resting-state FC (rsFC) strength within the posteromedial cortex (PMC), meaning the mean FC strength between PMC subregions, was differentially related to subsequent global neocortical amyloid depending on *APOE* genotype, with *APOE4* carriers showing more amyloid with declining rsFC within PMC but not *APOE4* non-carriers. **b)** The slope over time of mean FC strength during intentional encoding within medial temporal lobe (MTL) was differentially related to subsequent entorhinal tau depending on *APOE*, with *APOE4* carriers showing more subsequent tau with increasing encoding-FC within MTL but not *APOE4* non-carriers. **c)** The slope over time of mean FC strength during retrieval within the PMC was differentially related to subsequent entorhinal tau depending on *APOE*, with *APOE4* carriers showing more subsequent tau with increasing retrieval- FC within PMC, and *APOE4* non-carriers showing more subsequent tau with decreasing retrieval-FC within PMC. **d)** The slope over time of mean FC strength during retrieval between MTL and PMC was related to subsequent entorhinal tau, with more subsequent tau with increasing retrieval-FC between MTL and PMC. PET, Positron Emission Tomography; FC, functional connectivity; *APOE*, apolipoprotein E.

There was no significant association between global Aβ burden and the slope of FC strength within PMC during encoding or retrieval and no significant association between Aβ burden and the slope of FC within MTL or between MTL and PMC for any of the three paradigms (all *p*> 0.05; see Supplementary Tables S10 and S13 - 14).

Regarding entorhinal tau burden, we found an effect of encoding-FC within MTL depending on *APOE4* group (β=0.34 [95% CI 0.00, 0.68], t=1.980, *p*=0.0497; see Figure 2b and Supplementary Table S15). Specifically, *APOE4* carriers showed a steeper increasing slope of encoding-FC within MTL with more entorhinal tau (β=0.27 [95% CI -0.02, 0.55], t=1.876, *p*=0.066; see Supplementary Table S16), while *APOE4* non-carriers showed no association (β=-0.04 [95% CI -0.25, 0.17], t=-0.396, *p*=0.693; see Supplementary Table S17).

Across both *APOE4* groups, there was a main effect of retrieval-FC within PMC, with a steeper decreasing slope being related to more entorhinal tau (β=-0.35 [95% CI -0.54, -0.15], t=-3.540, *p*<0.001; see Supplementary Table S18). This effect was driven by *APOE4* non-carriers, as we also found a significant interaction between retrieval-FC within PMC and *APOE4* group (β=0.58 [95% CI 0.23, 0.94], t=3.259, *p*=0.001; see Figure 2c and Supplementary Table S18). *APOE4* carriers tended to show an increasing slope of retrieval-FC within PMC with more entorhinal tau (β=0.20 [95% CI -0.07, 0.47], t=1.459, *p*=0.151; see Supplementary Table S19), while *APOE4* non-carriers showed a decreasing slope of retrieval- FC within PMC with more entorhinal tau (β=-0.41 [95% CI -0.62, -0.20], t=-3.943, *p*<0.001; see Supplementary Table S20). Finally, there was a main effect of retrieval-FC between MTL and PMC, with an increasing slope being related to more entorhinal tau burden (β=0.28 [95% CI 0.06, 0.50], t=2.540, *p*=0.012; see Figure 2d and Supplementary Table S18).

In summary, we observed distinct associations between change in FC and subsequent AD pathology burden depending on *APOE4* genotype and fMRI paradigm (i.e. rest, encoding, and retrieval). In *APOE4* carriers, decreasing rsFC within PMC was related to more subsequent global Aβ burden, and increasing encoding-FC within MTL was related to more subsequent entorhinal tau burden. Decreasing retrieval-FC within PMC was related to more entorhinal tau, especially in *APOE4* non-carriers. Finally, increasing retrieval-FC between MTL and PMC was related to more entorhinal tau independent of *APOE4* status.

### Relationship between functional connectivity change and change in episodic memory performance

Finally, we investigated the association of the slopes of meta-ROI FC strength and the slopes of episodic memory performance using linear models including the interaction of FC slope by *APOE4* group, age, sex, and education. Episodic memory performance was measured via the Repeatable Battery for the Assessment of Neuropsychological Status (RBANS) delayed memory index score and the corrected hit rate from the object-location episodic memory fMRI retrieval session. With respect to changes in performance over time, we observed that the RBANS delayed memory index score increased over time (β=0.14 [95% CI 0.07, 0.21], t=4.199, *p*<0.001; see Supplementary Table S21), while the corrected hit rate decreased over time (β=-0.09 [95% CI -0.17, 0.01], t=-2.278, *p*=0.023; see Supplementary Table S22).

There was a main effect of encoding-FC within PMC for both measures of memory performance. Specifically, a decrease in PMC encoding-FC over time was related to an increase in the RBANS delayed memory index score (β=-0.27 [95% CI -0.50, -0.05], t=-2.412, *p*=0.017; see Supplementary Table S23) and the corrected hit rate (β=-0.37 [95% CI -0.59, -0.15], t=- 3.355, *p*=0.001; see Supplementary Table S24).

Further, there was a main effect of rsFC between MTL and PMC on the RBANS delayed memory index score, with an increase in FC being related to an increase in RBANS memory performance (β=0.31 [95% CI 0.02, 0.60], t=2.141, *p*=0.043; see Supplementary Table S25 and Figure 3a). Conversely, there was a main effect of encoding-FC between MTL and PMC, with a decrease in FC being related to an increase in the corrected hit rate performance (β=-0.26 [95% CI -0.47, 0.06], t=-2.519, *p*=0.013; see Supplementary Table S24 and Figure 3b).

**Figure 3.**
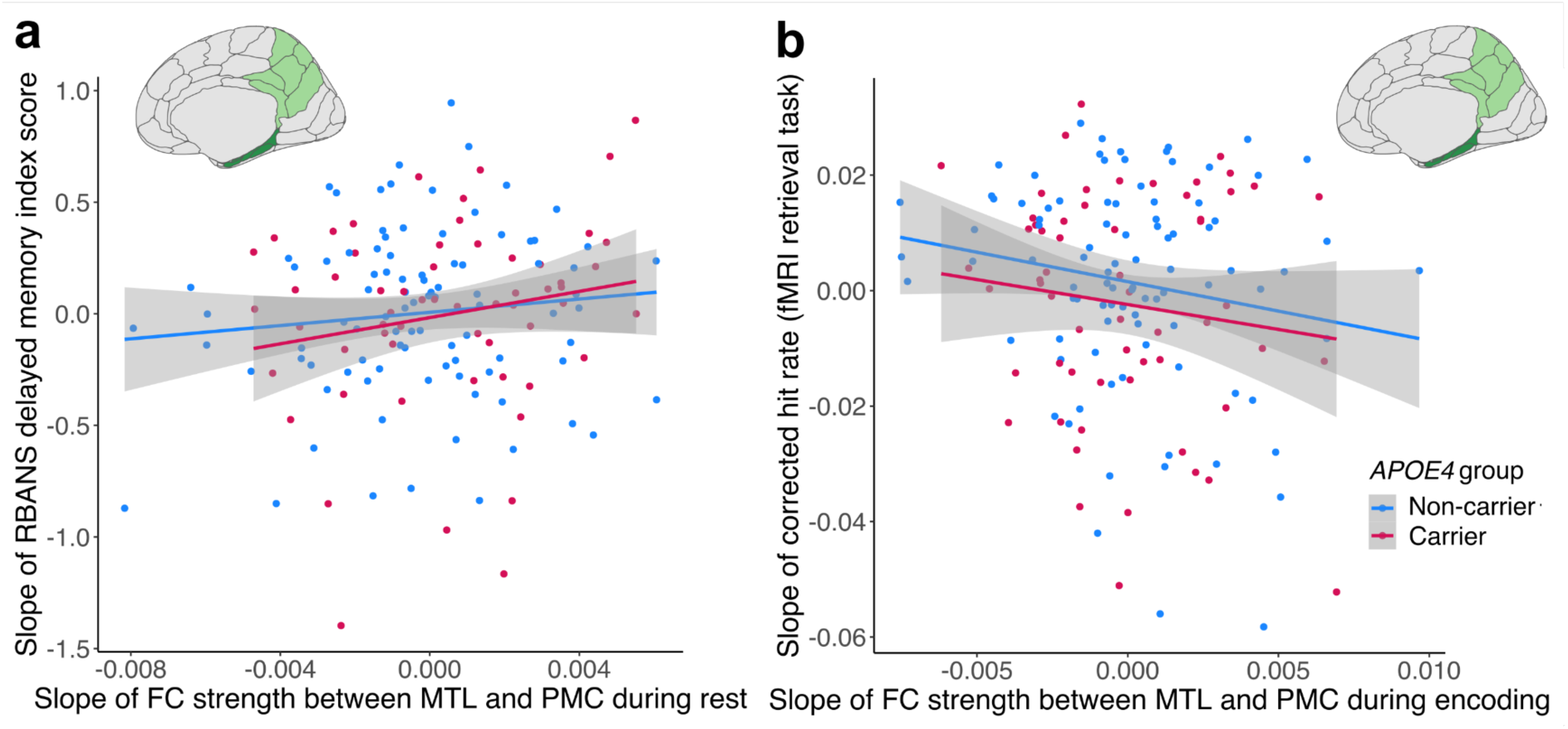
Episodic memory performance over time and the relationship with functional connectivity (FC) strength over time during rest and encoding. a) Increasing resting-state FC (rsFC) between MTL and PMC was related to an increasing RBANS delayed memory index score. The slope of cognitive performance was used for visualization. b) Increasing encoding- FC between MTL and PMC was related to decreasing encoding-task recognition performance. The slope of cognitive performance was used for visualization. RBANS, Repeatable Battery for the Assessment of Neuropsychological Status; FC, functional connectivity; fMRI, Functional Magnetic Resonance Imaging; *APOE*, apolipoprotein E.

There was no association between the slope of FC strength within MTL during any of the three paradigms and memory performance (all p>0.05), no effect of retrieval-FC strength for any region (all p>0.05), and no effects of *APOE4* genotype (all p>0.05). Further, there were no effects of the slope of FC strength on the slope of RBANS attention index score performance (all p>0.05). In summary, we observed differential associations between change in FC and change in episodic memory performance depending on the paradigm (i.e. rest, encoding, and retrieval) and region, with even opposing directions of associations for change in FC between MTL and PMC during rest versus during encoding (see Figure 3).

## Discussion

By analyzing longitudinal data across rest and task states, this study offers novel insights into FC changes within the episodic memory network and emphasizes differential associations with Alzheimer’s pathology and episodic memory in cognitively unimpaired older adults.

Our first aim was to study how changes in FC strength are linked to subsequent AD pathology burden. Within the PMC, decreasing FC was generally linked to more subsequent AD pathology, with decreasing FC at rest being associated with higher Aβ burden, and decreasing FC during retrieval being associated with higher tau burden. While prior studies reported higher PMC task activation and higher FC between the PMC and other brain regions with more Aβ^49–52^, there is also evidence for hypoconnectivity within the PMC with more Aβ^27,28,53^. The PMC is a highly connected hub region with the highest level of basal glucose energy consumption in the brain^54,55^. Beside hypoconnectivity, hypometabolism within the PMC has been linked to Aβ burden. For instance, Pascoal and colleagues showed that higher global Aβ burden was related to FDG-PET measured hypometabolism in the PMC and both interacted to predict subsequent cognitive decline^26^. PMC hypoconnectivity is less often reported in relation to tau pathology, however, lower FC strength within the default mode network (DMN), which includes the PMC, during rest was associated with more inferotemporal tau burden in cognitively unimpaired older adults^56^ and FDG-PET measured hypometabolism within the PMC was related to higher local tau burden in cognitively unimpaired^57^ and impaired^58^ individuals.

Importantly, we found that the *APOE4* genotype moderated the associations of FC and pathology, as reported in previous studies^39,59^. In our study, decreasing FC strength within the PMC during rest was associated with more Aβ in *APOE4* carriers, while the association of decreasing FC strength within the PMC during retrieval with more tau was driven by *APOE4* non-carriers. The *APOE4* genotype is more strongly related to Aβ^60–62^ than to tau pathology^63^, and *APOE4* carriers in our cohort showed significantly higher Aβ, but not tau burden. The *APOE4* genotype may predispose to vulnerability for greater activity-dependent pathology accumulation^15,64^. A recent study in the PREVENT-AD cohort hints towards this mechanism, showing that increasing PMC activity during retrieval was related to more subsequent global Aβ burden in *APOE4* carriers but not in non-carriers^65^. It is, however, unclear if there is a joint mechanism of simultaneously increasing PMC retrieval activity and decreasing FC within the PMC network, which may be pronounced in *APOE4* carriers. Further, our finding of decreasing retrieval-FC within PMC being associated with more entorhinal tau specifically in *APOE4* non-carriers in our sample should be investigated in future studies. This finding might be specific to primary age-related tauopathy (PART) in older individuals at low risk for AD^66,67^.

Overall, our results on longitudinal hypoconnectivity within the PMC suggest that disconnection or desynchronization among PMC subregions may be detrimental in the context of Aβ and tau pathology. This was previously reported for Aβ, we contribute novel evidence for this dynamic also for tau. Further, the *APOE4* genotype plays a critical but complex role in these associations that warrants further investigation.

Within the MTL, we report an association of increasing FC strength during encoding and more subsequent entorhinal tau burden in *APOE4* carriers. A recent study found higher FC during rest within the anterior-temporal network (including the perirhinal cortex and anterior hippocampus) with advancing age in a sample of cognitively unimpaired participants and impaired patients, however, the study did not have measures for tau specifically^68^. In non-demented *APOE4* carriers, increasing FC at rest within the hippocampus was reported in association with elevated plasma p-tau levels^39^. While prior research suggests that higher or increasing MTL activity during encoding is linked to more tau burden^69^, less is known about FC during encoding in relationship to tau.

Moreover, we found that increasing FC between the MTL and the PMC during retrieval was related to higher subsequent entorhinal tau burden. As FC between these regions is generally higher during retrieval and memory formation^70–72^, our findings suggest that hyperconnectivity specifically during task engagement may be related to detrimental pathological processes. Prior resting-state studies reported higher FC between the hippocampus and the retrosplenial cortex with more medial-parietal tau burden^25^ and higher FC strength seems to accelerate tau spread from the MTL to the PMC^23,24^. Higher or increasing retrieval activity of the MTL and the PMC was further reported in association with more Aβ pathology^65,73,74^, but studies on the association with tau and investigating FC during retrieval are lacking. Interestingly, in *APOE4* carriers, we observed further associations of increasing FC in task-engaged regions and higher tau pathology, i.e. within the MTL during encoding and within the PMC during retrieval. This supports the theory that hyperconnectivity particularly of task-engaged regions is detrimental regarding pathology, potentially especially pronounced in *APOE4* carriers who are at higher risk for AD. The underlying mechanism, e.g. heightened tau spread and accumulation specifically under these circumstances, needs to be determined using study designs including longitudinal tau-PET.

Our second aim was to investigate the association of change in FC strength in episodic memory regions with change in episodic memory performance. It is still unclear whether increasing FC strength represents beneficial or compensatory processes to maintain cognitive performance when facing pathology or whether it is part of a vicious cycle of pathology accumulation and spread^14,75,76^. In our study, we found that decreasing FC in the PMC during encoding was associated with increasing memory performance in both the RBANS delayed memory index score and memory performance in the fMRI object-location episodic memory task. Notably, PMC regions usually deactivate during memory encoding^77,78^. This hints towards maintaining low FC strength within this task-negative region as being beneficial with respect to cognitive changes, whereas increasing FC in the PMC during encoding might be detrimental. While little is known about FC during memory tasks, prior studies showed opposite relationships for PMC connectivity at rest and memory performance. Specifically, reduced FC strength within the PMC during rest at baseline was related to longitudinal decrease in episodic memory^27,30^. These rsFC studies hint towards beneficial effects of high network connectivity during rest, where the PMC is typically engaged in healthy adults^54^.

Maintaining low FC strength between the MTL and the PMC during encoding seemed to be also beneficial for episodic memory performance, similar to FC within PMC during encoding, as we found that increasing within and between network FC during encoding was related to a declining corrected hit rate. More research has been conducted using FC at rest, with prior studies^9,33,34,79^ reporting higher rsFC strength between MTL and PMC in cognitively unimpaired older adults as being beneficial for episodic memory, consistent with our rsFC results.

Our results add to the debate on beneficial and detrimental effect of FC on cognition and underscore the importance of taking into account which specific brain regions or networks are involved and whether connectivity is measured during rest, task encoding or task retrieval.

Our third aim was to investigate whether similar dynamics of connectivity change occur during resting state as during episodic memory task paradigms. While change in FC at rest was related to Aβ burden, change in FC during components of the episodic memory task, i.e. encoding and retrieval, was related to tau burden. This observation provides an interesting starting point for future studies to investigate data from multiple fMRI paradigms in relation to AD pathology^15^. Furthermore, we found differential implications for episodic memory performance of increasing or decreasing FC between the MTL and the PMC depending on the fMRI paradigm or stage of memory processing. Our findings point to opposing directions of FC changes during rest versus encoding in relation to cognitive trajectories. Interestingly, FC changes between MTL and PMC during rest versus encoding were inversely related, in contrast to the otherwise positive correlations observed between FC slopes.

Studies combining rest and task fMRI are rare, however, Cassady and colleagues reported less similarity in FC of the episodic memory network between rest and an object-scene mnemonic discrimination task in cognitively unimpaired older compared to younger adults, especially with more local PET-measured tau, and less similarity in FC was related to worse object discrimination^11^. Kizilirmak and colleagues investigated voxelwise DMN resting-state mean percent amplitude of fluctuation (mPerAF) and DMN fMRI activation during successful memory encoding in cognitively unimpaired younger and older adults. Cognitively unimpaired older adults showed less encoding-related deactivation and lower resting-state amplitudes in the DMN. Older adults with more DMN deactivation during encoding and higher DMN resting-state amplitudes showed higher performance in an independent episodic memory test, while this was not the case for younger adults^80^. Together, these two studies suggest that, in older adults, maintaining a stable network architecture across rest and task states, while retaining the flexibility to modulate networks in response to task demands, may be beneficial for cognition.

There are several limitations of the present study that should be considered. First, while we investigated the slope of mean FC strength over time within three meta-ROIs (within MTL, within PMC, and between MTL and PMC), we cannot provide specificity at the level of individual ROIs, like the anterior hippocampus. While longitudinal ROI-to-ROI group-level analysis would have provided this specificity, this method requires complete datasets. Thus, our chosen statistical approach offered the advantage of including a larger sample with data over up to four years, as linear mixed models enable the inclusion of participants with missing time points. Furthermore, our methods were preregistered, strengthening the transparency and reproducibility of our approach. Future studies could address these limitations by investigating datasets with complete longitudinal fMRI data or by employing more advanced methods to investigate effective connectivity^81^ and brain states^82,83^. Second, we had no longitudinal PET measurements available, thus, we cannot comment on the interplay of change in FC strength and accumulation rates or patterns of AD pathology. Future multimodal studies could include follow-up PET sessions to address those interactions. Third, we did not relate changes in FC strength to changes in fMRI task activity. Future multimodal studies could address complex network disruptions, e.g. whether decreasing memory-task FC within the PMC is occurring simultaneously with increasing memory-task activity in the precuneus^65,80^, particularly in the context of AD pathology. Fourth, our sample consisted of highly-educated older adults and sex was not balanced. Future research could aim at including participants from more diverse backgrounds and avoid biases in sex distribution and further variables.

To conclude, we provide support that longitudinal changes in FC strength during rest and task are differentially related to AD pathology and episodic memory in cognitively unimpaired older adults. Hypoconnectivity in the PMC over time was related to higher subsequent global neocortical Aβ and entorhinal tau burden, while hyperconnectivity involving the MTL was related to higher subsequent entorhinal tau burden. As some changes in FC strength interacted with the *APOE4* genotype, complex mechanisms seem to be at play. Further, increasing FC strength between the MTL and the PMC during rest, contrary to FC strength during encoding, was beneficial for episodic memory performance. Notably, changes in FC strength at rest were associated with Aβ burden, while changes in FC strength during memory encoding and retrieval were associated with tau burden. These results underscore that pathology-related aberrant network changes present differently across rest and task states, with distinct effects on episodic memory.

## Methods

### Participants

One hundred fifty-two cognitively unimpaired older adults from the PREVENT-AD cohort were included in this study^45,46^. PREVENT-AD is a longitudinal Canadian cohort study that started in 2011 and includes participants over the age of 60 at the time of enrollment. Participants need to have a self-reported parental history of AD-like dementia or two siblings with AD-like dementia. Individuals aged 55 to 59 were also included if their age was within 15 years of their affected relative’s age of disease onset. No participants had major psychiatric or neurological conditions at baseline and all participants were assessed for normal cognition with the Montreal Cognitive Assessment (MoCA)^84^ and the Clinical Dementia Rating (CDR) scale^85^. If concerns about cognition were raised (MoCA score below 27 or CDR score above 0), participants were required to undergo comprehensive neuropsychological testing, and those presenting with probable mild cognitive impairment (MCI) were excluded. The data utilized in this study were drawn from data release 7.0, with parts of the dataset being publicly accessible (https://openpreventad.loris.ca/) and parts of the dataset being accessible to registered researchers only (https://registeredpreventad.loris.ca/). Follow-up fMRI and cognitive testing sessions were performed over 48 months in a subset of participants. Specifically, participants completed a 12-month, 24-month, and 48-month follow-up after baseline. For an overview of the included sessions, see Figure 1a.

One hundred sixty-five participants had at least baseline resting-state, baseline task fMRI, and PET data available. However, 7 participants were excluded because of incomplete fMRI data, 3 participants were excluded after quality control of task-fMRI data, and another 3 participants were excluded after quality control of resting-state fMRI data. Additionally, 10 sessions of resting-state imaging were excluded after quality control, among them one baseline session. These exclusions based on fMRI quality control were applied when more than 20% of volumes were flagged as outliers due to excessive motion (see “MRI processing” for details).

All study procedures and experimental protocols were approved by the McGill University Institutional Review Board and/or the Douglas Mental Health Institute Research Ethics Board in accordance with the Declaration of Helsinki. Written informed consent was obtained from all participants prior to each experimental procedure and they were financially compensated for their time.

### *APOE* genotyping

The participants were genotyped for *APOE* using a QIASymphony apparatus, details are described by Tremblay-Mercier et al.^46^. Those participants carrying at least one *APOE4* allele were assigned to the carrier group, while those without an *APOE4* allele were placed in the non-carrier group.

## Assessment of episodic memory performance

For cognitive testing, the RBANS was used at each session^86^. To minimize practice effects, different versions were used at each follow-up session. Our primary variable of interest was the RBANS delayed memory index score, which combines word-list recognition with delayed recall of figures, stories, and word lists. To assess the specificity of the relationships between longitudinal memory performance and FC, we used the attention index score as a memory-independent measure in a control analysis. The attention index score is based on two digit-span tests. Two participants had missing baseline data for cognitive testing, therefore, we used the RBANS data measured three months after baseline in those cases.

Participants performed an object-location episodic memory task in the fMRI scanner that consisted of an encoding session and a retrieval session 20 minutes later^48,65^. In the encoding session, participants were presented with 48 visual stimuli (colored line drawings of everyday objects) that appeared on either the right or left side of the screen. Participants were instructed to press a button to indicate on which side the stimulus appeared. During the retrieval session, participants were presented with 48 previously seen objects (from the encoding phase) and 48 new objects. Participants made a forced-choice response regarding recognition using four options: i) “The object is FAMILIAR, but you don’t remember the location” (“F”); ii) “You remember the object and it was previously on the LEFT” (“L”); iii) “You remember the object and it was previously on the RIGHT” (“R”); and iv) “The object is NEW” (“N”) (see Supplementary Figure S1). Details are described by Rabipour and colleagues^48^. As a second measure of episodic memory performance beside the RBANS delayed memory index score, we used the corrected hit rate from the described retrieval task. The corrected hit rate was specified as hits (i.e. responses “Familiar”, ”Remember-Left” or ”Remember-Right” to previously shown objects) minus false alarms (responses “Familiar”, ”Remember-Left” or ”Remember-Right” to novel objects).

## MRI acquisition and processing

All MRI data were acquired using a Siemens Tim Trio 3-Tesla MRI scanner (Siemens Medical Solutions, Erlangen, Germany) at the Cerebral Imaging Centre, Douglas Mental Health University Institute, with a Siemens standard 12- or 32-channel coil^46^.

T1-weighted anatomical MPRAGE images were obtained (TR = 2300 ms, TE = 2.98 ms, TI = 900 ms, flip angle = 9°, FOV = 256x240x176) and had a 1 mm isotropic voxel resolution. Resting-state fMRI data were collected over 10 minutes using an EPI sequence (TR = 2000 ms, TE = 30 ms, flip angle = 90°, FOV = 256x256 mm, 32 slices) with a 4 mm isotropic voxel resolution. For task-fMRI data, an EPI sequence (TR = 2000 ms, TE = 30 ms, flip angle = 90°, FOV = 256x256 mm, 32 slices) with a 4 mm isotropic voxel resolution was used as well. During each task-fMRI session, participants completed encoding for 6 minutes and retrieval for 15 minutes of the described object-location episodic memory task (see “Assessment of episodic memory performance”). Additionally, fieldmaps were acquired to correct for spatial distortions caused by field inhomogeneities during unwarping.

Functional and structural data were preprocessed using MATLAB and SPM version 12^87^ as well as the CONN toolbox, version 22.a^88^. Functional data were slice-time corrected, realigned, and co-registered to the structural T1-weighted image of the respective session. The structural images were segmented into distinct tissue types (gray matter, white matter, and cerebrospinal fluid). All MRI data were normalized to the Montreal Neurological Institute (MNI) space, utilizing the IXI549Space template implemented in SPM12. We defined in our preregistration^44^ to not apply smoothing, as we aimed at keeping the task-preprocessing pipeline similar to the resting-state pipeline, where typically no smoothing is applied for Roi-to-Roi analyses.

Outliers caused by excessive head motion were identified using ART^89^, with volumes flagged if framewise displacement exceeded 0.5 mm/TR or if the global intensity z-score was greater than 3. Sessions with more than 20% of volumes flagged as outliers were removed from further analyses (see “Participants”). During denoising of functional data, confounding effects were regressed out, including realignment parameters and their first-order derivatives, flagged outlier volumes, cerebral white matter, and cerebrospinal fluid signal derived via an anatomical component-based noise correction procedure (CompCor)^90^. A band-pass filter (0.008 Hz to 0.09 Hz) was applied to reduce noise from physiological and motion artifacts^91^.

ROIs for ROI-to-ROI functional connectivity analysis were selected a-priori based on literature research on episodic memory brain areas. They included (i) the MTL regions anterior and posterior hippocampus, lateral and medial parahippocampal cortex, entorhinal cortex, and anterior and posterior perirhinal cortex, and ii) the PMC regions precuneus (consisting of a superior, inferior, medial, and lateral ROI), posterior cingulate cortex, and retrosplenial cortex. The ROIs were derived using the Brainnetome atlas^47^.

The whole resting-state, encoding, and retrieval sessions were used to assess FC strength. The volumes per session were weighted equally and convolved with a canonical hemodynamic response function to reduce the influence of the first volumes. BOLD time-series were extracted from the 13 predefined Brainnetome ROIs and used for ROI-to-ROI first-level analyses. This generated a functional connectivity matrix for each participant and session, representing Fisher’s r-to-z transformed correlation coefficients between each pair of ROIs. We then calculated the mean FC strength for each participant and session for three meta-ROIs, i.e. (i) all ROIs within MTL, ii) all ROIs within PMC, and iii) between MTL and PMC ROIs.

## PET acquisition and processing

PET scanning was conducted at the McConnell Brain Imaging Centre at the Montreal Neurological Institute (Quebec, Canada), using a brain-dedicated Siemens/CTI high-resolution research tomograph. The procedures for data acquisition and processing followed the methods outlined by Yakoub and colleagues^92^. Briefly, Aβ-PET images using NAV were acquired 40– 70 minutes post-injection (∼6 mCi dose), and tau-PET images using FTP were acquired 80– 100 minutes post-injection (∼10 mCi dose). Five-minute frames and an attenuation scan were additionally obtained. PET images were reconstructed using a 3D OP-OSEM algorithm with 10 iterations and 16 subsets, applying decay and motion corrections, as well as 3D scatter estimation for scatter correction. T1-weighted MRI images were segmented based on the Desikan-Killiany atlas using FreeSurfer version 5.3. PET images were realigned, temporally averaged, co-registered to the T1-weighted MRI image (from the scan closest in time), masked to exclude cerebrospinal fluid (CSF), and smoothed with a 6 mm Gaussian kernel. Standardized uptake value ratios (SUVRs) were calculated for each Freesurfer ROI, comparing tracer uptake to cerebellar gray matter for Aβ-PET and inferior cerebellar gray matter for tau-PET. The entire PET dataset was processed using a standard pipeline (https://github.com/villeneuvelab/vlpp). For tau, a region-of-interest (ROI) approach was used, specifically evaluating bilateral entorhinal FTP SUVR by averaging uptake in both entorhinal cortices. For Aβ, global neocortical NAV SUVR consisting of bilateral lateral and medial frontal, parietal, and lateral temporal regions was investigated. Aβ and tau pathology values were non-normally distributed, therefore, a Box-Cox transformation was applied to the PET data to achieve a closer approximation of a normal distribution. Biomarker status was determined based on research cut-offs established by the PREVENT-AD Research Group, with entorhinal FTP SUVR > 1.3 and global neocortical NAV SUVR > 1.39 indicating biomarker positivity.

## Statistical analyses

The statistical analyses were conducted using R^93^, version 4.1.2, implemented within RStudio^94^. The R code used for analyses is publicly available (https://github.com/fislarissa/MTL_PMC_rs_and_task_FC). For linear models, it was ensured that the assumption of homoscedasticity was met using the Breusch-Pagan test. When heteroscedasticity was detected, robust standard errors are reported. We ensured that no multicollinearity was present with a variance inflation factor (VIF) below 5. Furthermore, the assumption of a normal distribution of residuals was assessed using quantile-quantile (QQ) plots for each model. A significance threshold of *p* < 0.05 was used. For exploratory analyses, multiple comparisons were controlled using False-discovery-rate (FDR) correction according to the Benjamini-Hochberg procedure. All preregistered models were analyzed as confirmatory and hypothesis-driven, and no correction for multiple comparisons across these models was applied^43,44^. For linear mixed models, a random intercept and slope for participant was included. If this led to a singular fit, only a random intercept was used. The covariates age at baseline, sex, and education were used in all models.

First, we investigated the influence of time and the interaction of time by *APOE4* group on longitudinal FC strength in linear mixed models separately for each meta-ROI and paradigm (e.g. *FC strength within MTL at rest ∼ Time * APOE4 group + Age + Sex + Education)*. Exploratively, we compared the slopes of FC strength between paradigms using correlation analyses.

Second, we investigated the effects of the slope of FC over time and the interaction of slope of FC by *APOE4* group on subsequent Aβ and tau PET burden, including time between the baseline MRI and PET session as additional covariate (e.g. Aβ*-PET burden ∼ Slope of FC strength within MTL at rest * APOE4 group + Slope of FC strength within PMC at rest * APOE4 group + Slope of FC strength between MTL and PMC at rest * APOE4 group + Age + Sex + Education + Time between baseline MRI and PET)*.

Third, we investigated the influence of time and *APOE4* group on the RBANS delayed memory index score and corrected hit rate from the fMRI task over time (e.g. *RBANS delayed memory index score ∼ Time + APOE4 group + Age + Sex + Education)*. We then investigated the effects of the slope of FC over time and the interaction of slope of FC by *APOE4* group on slope of memory performance (e.g. *Slope of RBANS delayed memory index score ∼ Slope of FC strength within MTL at rest * APOE4 group + Slope of FC strength within PMC at rest * APOE4 group + Slope of FC strength between MTL and PMC at rest * APOE4 group + Age + Sex + Education)*. As a control analysis, we investigated the RBANS attention score instead of the measures of episodic memory performance, where we expected no associations with change in FC strength in the memory-related brain areas we investigated. The models investigating AD pathology and cognition were conducted separately for the resting-state, encoding, and retrieval timeseries, respectively, to avoid multicollinearity.

## Acknowledgements

We want to thank the participants of the PREVENT-AD study for their time and effort as well as the researchers involved in building up the cohort https://preventad.loris.ca/acknowledgements/acknowledgements.php?DR=7.0. This work was supported by the German Research Foundation (DFG; Project-ID 425899996, CRC1436 to A.M and E.N.M; Project-ID 362321501, RTG 2413 to A.M. and L.F.).

## Author contributions

Conceptualization: LF, JNA, ENM, AM. Methodology: LF, JNA, AM. Formal analysis: LF. Data Acquisition: JTM, JR, APB, MNR, SV, PREVENT-AD Research Group. Image processing: JR, LF. Visualization: LF. Image analysis and modelling: LF, AM. Investigation: LF, AM. Supervision: JNA, AM. Project administration: AM, SV. Funding acquisition: AM. Resources: AM, SV, PREVENT-AD Research Group. Writing original draft preparation: LF, AM. Writing – review and editing: All authors. All authors read and approved the final manuscript.

## Data availability statement

Some of the data used in this manuscript is openly available at https://openpreventad.loris.ca. The remaining data can be shared upon approval by the scientific committee at the Centre for Studies on Prevention of Alzheimer’s Disease (StoP-AD) at the Douglas Mental Health University Institute after registration at https://registeredpreventad.loris.ca. Code for PET preprocessing is available at https://github.com/villeneuvelab/vlpp and code for statistical analysis is available at https://github.com/fislarissa/MTL_PMC_rs_and_task_FC.

