## Supplementary Material for "Longitudinal functional connectivity during rest and task is differentially related to Alzheimer’s pathology and episodic memory in older adults"

-

### Supplementary material

Larissa Fischer[1,2]*, Jenna N. Adams[2], Eóin N. Molloy[1,3], Jennifer Tremblay-Mercier[4], Jordana Remz[4], Alexa Pichet Binette[5,6,7], M. Natasha Rajah[4,8,9], Sylvia Villeneuve[4,8], Anne Maass[1,10], PREVENT-AD Research Group

[1] German Center for Neurodegenerative Diseases (DZNE), Magdeburg, Germany
[2] Department of Neurobiology and Behavior, University of California, Irvine, USA
[3] Department of Radiology & Nuclear Medicine, Faculty of Medicine, Otto von Guericke University, Magdeburg, Germany
[4] StoP-AD Centre, Douglas Mental Health Institute Research Centre, Montréal, Canada
[5] Department of Physiology and Pharmacology, Université de Montréal, Montréal, Canada
[6] Centre de Recherche de l'Institut Universitaire de Gériatrie de Montréal, Montréal, Canada
[7] Clinical Memory Research Unit, Department of Clinical Sciences Malmö, Lund University, Lund, Sweden
[8] Department of Psychiatry, McGill University, Montréal, Canada
[9] Department of Psychology, Toronto Metropolitan University, Toronto, Canada
[10] Institute for Biology, Otto von Guericke University, Magdeburg, Germany

***Corresponding author:**
Larissa Fischer

Leipziger Straße 44
39106 Magdeburg
Germany

A complete listing of the PREVENT-AD Research Group can be found at: https://preventad.loris.ca/acknowledgements/acknowledgements.php?DR=7.0&authors. Data used in preparation of this article were obtained from the Pre-symptomatic Evaluation of Experimental or Novel Treatments for Alzheimer's Disease (PREVENT-AD) program (https://www.centre-stopad.com/en/).

**Keywords**

Aging, Alzheimer's disease, fMRI, Functional Connectivity, Episodic Memory, *APOE4*

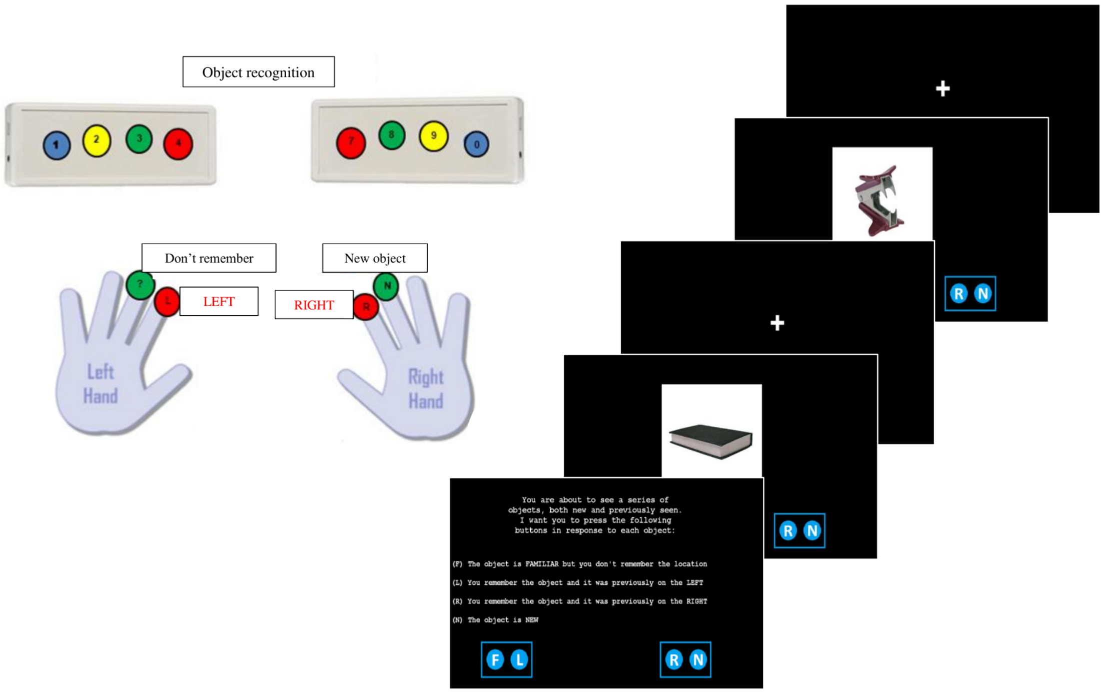


**Supplementary Fig. 1. Episodic memory retrieval task paradigm:** After the encoding session, participants were presented with 48 “old” items (previously shown at encoding) and 48 “new” (not previously seen) items. The items were colored object stimuli presented in the center of the computer screen. Participants completed a 4-alternative forced choice task: “Familiar” (F) to indicate that the object is familiar, but the participant cannot remember whether it was on the left or right, “Remember-Left” (L) to indicate that the participant remembered the object and that it was on the left, “Remember-Right” (R) to indicate that the participant remembered the object and that it was on the right, and “New” (N) to indicate that the object was not previously presented. The presentation rate was 4 seconds. Figure adapted from[65].

#### Supplementary Tab. 1. Linear model of effects of covariates on rsFC within MTL over time

| | Resting-state functional connectivity strength within MTL | | | | | | | | | |
|---|---|---|---|---|---|---|---|---|---|---|
| *Predictors* | *Estimates* | *std. Error* | *std. Beta* | *standardized std. Error* | *CI* | *standardized CI* | *Statistic* | *std. Statistic* | *p* | *std. p* |
| (Intercept) | 0.26 | 0.08 | 0.49 | 0.12 | 0.10 – 0.42 | 0.26 – 0.72 | 3.17 | 4.18 | **0.002** | **<0.001** |
| Time in days | -0.01 | 0.00 | -0.10 | 0.04 | -0.02 – -0.00 | -0.19 – -0.01 | -2.20 | -2.20 | **0.029** | **0.029** |
| *APOE4* Group [carrier] | -0.02 | 0.01 | -0.20 | 0.12 | -0.04 – 0.01 | -0.44 – 0.04 | -1.37 | -1.63 | 0.172 | 0.105 |
| Age in years | -0.00 | 0.00 | -0.02 | 0.06 | -0.00 – 0.00 | -0.14 – 0.10 | -0.37 | -0.37 | 0.715 | 0.715 |
| Sex [female] | -0.05 | 0.01 | -0.61 | 0.13 | -0.08 – -0.03 | -0.87 – -0.36 | -4.78 | -4.78 | **<0.001** | **<0.001** |
| Education in years | 0.00 | 0.00 | 0.06 | 0.06 | -0.00 – 0.00 | -0.06 – 0.18 | 0.94 | 0.94 | 0.348 | 0.348 |

| | | | | | | | | | | |
|---|---|---|---|---|---|---|---|---|---|---|
| Time in days × *APOE4* Group [carrier] | 0.00 | 0.01 | 0.04 | 0.08 | -0.01 – 0.02 | -0.12 – 0.19 | 0.49 | 0.49 | 0.625 | 0.625 |
| **Random Effects** | | | | | | | | | | |
| $\sigma^2$ | 0.00 | | | | | | | | | |
| $\tau_{00\ subj}$ | 0.00 | | | | | | | | | |
| ICC | 0.36 | | | | | | | | | |
| $N_{subj}$ | 152 | | | | | | | | | |
| Observations | 469 | | | | | | | | | |
| Marginal $R^2$ / Conditional $R^2$ | 0.099 / 0.424 | | | | | | | | | |

rsFC = resting-state functional connectivity. MTL = medial temporal lobe. CI = 95% confidence interval.

#### Supplementary Tab. 2. Linear model of effects of covariates on rsFC within PMC over time

| | **Resting-state functional connectivity strength within PMC** | | | | | | | | | |
|---|---|---|---|---|---|---|---|---|---|---|
| *Predictors* | *Estimates* | *std. Error* | *std. Beta* | *standardized std. Error* | *CI* | *standardized CI* | *Statistic* | *std. Statistic* | *p* | *std. p* |
| (Intercept) | 0.42 | 0.11 | 0.34 | 0.12 | 0.21 – 0.64 | 0.10 – 0.58 | 3.87 | 2.84 | **<0.001** | **0.005** |
| Time in days | -0.02 | 0.01 | -0.13 | 0.04 | -0.03 – -0.01 | -0.21 – -0.04 | -2.90 | -2.90 | **0.004** | **0.004** |
| *APOE4* Group [carrier] | 0.01 | 0.02 | 0.06 | 0.13 | -0.02 – 0.04 | -0.19 – 0.31 | 0.74 | 0.47 | 0.460 | 0.639 |
| Age in years | 0.00 | 0.00 | 0.03 | 0.06 | -0.00 – 0.00 | -0.10 – 0.15 | 0.41 | 0.41 | 0.685 | 0.685 |
| Sex [female] | -0.06 | 0.01 | -0.56 | 0.13 | -0.09 – -0.03 | -0.82 – -0.30 | -4.22 | -4.22 | **<0.001** | **<0.001** |
| Education in years | -0.00 | 0.00 | -0.02 | 0.06 | -0.00 – 0.00 | -0.14 – 0.10 | -0.31 | -0.31 | 0.759 | 0.759 |
| Time in days × *APOE4* Group [carrier] | 0.01 | 0.01 | 0.08 | 0.08 | -0.01 – 0.03 | -0.07 – 0.23 | 1.04 | 1.04 | 0.300 | 0.300 |
| **Random Effects** | | | | | | | | | | |
| $\sigma^2$ | 0.01 | | | | | | | | | |
| $\tau_{00\ subj}$ | 0.00 | | | | | | | | | |
| ICC | 0.41 | | | | | | | | | |
| $N_{subj}$ | 152 | | | | | | | | | |
| Observations | 469 | | | | | | | | | |

| | |
|---|---|
| Marginal $R^2$ / Conditional $R^2$ | 0.084 / 0.460 |

rsFC = resting-state functional connectivity. PMC = posteromedial cortex. CI = 95% confidence interval.

#### Supplementary Tab. 3. Linear model of effects of covariates on rsFC between MTL and PMC over time

| | Resting-state functional connectivity strength between MTL and PMC | | | | | | | | | |
|---|---|---|---|---|---|---|---|---|---|---|
| *Predictors* | *Estimates* | *std. Error* | *std. Beta* | *standardized std. Error* | *CI* | *standardized CI* | *Statistic* | *std. Statistic* | *p* | *std. p* |
| (Intercept) | 0.08 | 0.07 | 0.37 | 0.12 | -0.06 – 0.22 | 0.13 – 0.60 | 1.07 | 3.10 | 0.287 | **0.002** |
| Time in days | -0.01 | 0.00 | -0.09 | 0.05 | -0.02 – -0.00 | -0.18 – -0.00 | -1.99 | -1.99 | **0.048** | **0.048** |
| *APOE4* Group [carrier] | -0.01 | 0.01 | -0.16 | 0.12 | -0.03 – 0.01 | -0.41 – 0.08 | -0.73 | -1.31 | 0.466 | 0.191 |
| Age in years | -0.00 | 0.00 | -0.00 | 0.06 | -0.00 – 0.00 | -0.12 – 0.12 | -0.03 | -0.03 | 0.976 | 0.976 |
| Sex [female] | -0.04 | 0.01 | -0.47 | 0.13 | -0.05 – -0.02 | -0.73 – -0.22 | -3.64 | -3.64 | **<0.001** | **<0.001** |
| Education in years | 0.00 | 0.00 | 0.07 | 0.06 | -0.00 – 0.00 | -0.05 – 0.19 | 1.10 | 1.10 | 0.272 | 0.272 |
| Time in days × *APOE4* Group [carrier] | 0.01 | 0.01 | 0.13 | 0.08 | -0.00 – 0.02 | -0.03 – 0.29 | 1.59 | 1.59 | 0.112 | 0.112 |
| **Random Effects** | | | | | | | | | | |
| $\sigma^2$ | 0.00 | | | | | | | | | |
| $\tau_{00\ subj}$ | 0.00 | | | | | | | | | |
| ICC | 0.35 | | | | | | | | | |
| $N_{subj}$ | 152 | | | | | | | | | |
| Observations | 469 | | | | | | | | | |
| Marginal $R^2$ / Conditional $R^2$ | 0.066 / 0.391 | | | | | | | | | |

rsFC = resting-state functional connectivity. MTL = medial temporal lobe. PMC = posteromedial cortex. CI = 95% confidence interval.

#### Supplementary Tab. 4. Linear model of effects of covariates on encoding-FC within MTL over time

| | Functional connectivity strength during encoding within MTL | | | | | | | | | |
|---|---|---|---|---|---|---|---|---|---|---|
| *Predictors* | *Estimates* | *std. Error* | *std. Beta* | *standardized std. Error* | *CI* | *standardized CI* | *Statistic* | *std. Statistic* | *p* | *std. p* |
| (Intercept) | 0.44 | 0.09 | 0.21 | 0.12 | 0.25 – 0.62 | -0.04 – 0.46 | 4.68 | 1.69 | **<0.001** | 0.093 |

| | | | | | | | | | | |
|---|---|---|---|---|---|---|---|---|---|---|
| Time in days | 0.01 | 0.00 | 0.05 | 0.04 | -0.00 – 0.01 | -0.03 – 0.14 | 1.20 | 1.20 | 0.232 | 0.232 |
| *APOE4* Group [carrier] | 0.01 | 0.01 | 0.09 | 0.13 | -0.02 – 0.03 | -0.17 – 0.35 | 0.55 | 0.68 | 0.581 | 0.500 |
| Age in years | -0.00 | 0.00 | -0.13 | 0.07 | -0.01 – 0.00 | -0.26 – 0.00 | -1.97 | -1.97 | 0.051 | 0.051 |
| Sex [female] | -0.03 | 0.01 | -0.36 | 0.14 | -0.06 – -0.01 | -0.63 – -0.09 | -2.62 | -2.62 | **0.010** | **0.010** |
| Education in years | 0.00 | 0.00 | 0.00 | 0.06 | -0.00 – 0.00 | -0.13 – 0.13 | 0.01 | 0.01 | 0.995 | 0.995 |
| Time in days × *APOE4* Group [carrier] | -0.00 | 0.01 | -0.02 | 0.08 | -0.02 – 0.01 | -0.17 – 0.13 | -0.26 | -0.26 | 0.793 | 0.793 |
| **Random Effects** | | | | | | | | | | |
| $\sigma^2$ | 0.00 | | | | | | | | | |
| $\tau_{00\ subj}$ | 0.00 | | | | | | | | | |
| ICC | 0.43 | | | | | | | | | |
| $N_{subj}$ | 152 | | | | | | | | | |
| Observations | 457 | | | | | | | | | |
| Marginal $R^2$ / Conditional $R^2$ | 0.044 / 0.454 | | | | | | | | | |

FC = functional connectivity. MTL = medial temporal lobe. CI = 95% confidence interval.

#### Supplementary Tab. 5. Linear model of effects of covariates on encoding-FC within PMC over time

| | Functional connectivity strength during encoding within PMC | | | | | | | | | |
|---|---|---|---|---|---|---|---|---|---|---|
| *Predictors* | *Estimates* | *std. Error* | *std. Beta* | *standardized std. Error* | *CI* | *standardized CI* | *Statistic* | *std. Statistic* | *p* | *std. p* |
| (Intercept) | 0.35 | 0.12 | 0.25 | 0.12 | 0.12 – 0.58 | 0.01 – 0.49 | 2.98 | 2.07 | **0.003** | **0.040** |
| Time in days | 0.00 | 0.01 | 0.02 | 0.05 | -0.01 – 0.01 | -0.07 – 0.11 | 0.43 | 0.43 | 0.668 | 0.668 |
| *APOE4* Group [carrier] | 0.00 | 0.02 | -0.03 | 0.13 | -0.03 – 0.03 | -0.29 – 0.22 | 0.10 | -0.24 | 0.918 | 0.812 |
| Age in years | 0.00 | 0.00 | 0.11 | 0.06 | -0.00 – 0.01 | -0.02 – 0.24 | 1.70 | 1.70 | 0.092 | 0.092 |
| Sex [female] | -0.04 | 0.02 | -0.36 | 0.13 | -0.07 – -0.01 | -0.62 – -0.09 | -2.68 | -2.68 | **0.008** | **0.008** |
| Education in years | 0.00 | 0.00 | 0.01 | 0.06 | -0.00 – 0.00 | -0.11 – 0.14 | 0.20 | 0.20 | 0.839 | 0.839 |
| Time in days × *APOE4* Group [carrier] | 0.01 | 0.01 | 0.08 | 0.08 | -0.01 – 0.03 | -0.07 – 0.24 | 1.05 | 1.05 | 0.295 | 0.295 |

**Random Effects**

| | |
|---|---|
| $\sigma^2$ | 0.01 |
| $\tau_{00\ subj}$ | 0.01 |
| ICC | 0.39 |
| $N_{subj}$ | 152 |
| Observations | 457 |
| Marginal $R^2$ / Conditional $R^2$ | 0.053 / 0.427 |

FC = functional connectivity. PMC = posteromedial cortex. CI = 95% confidence interval.

#### Supplementary Tab. 6. Linear model of effects of covariates on encoding-FC between MTL and PMC over time

| | Functional connectivity strength during encoding between MTL and PMC | | | | | | | | | |
|---|---|---|---|---|---|---|---|---|---|---|
| *Predictors* | *Estimates* | *std. Error* | *std. Beta* | *standardized std. Error* | *CI* | *standardized CI* | *Statistic* | *std. Statistic* | *p* | *std. p* |
| (Intercept) | -0.01 | 0.07 | 0.10 | 0.11 | -0.15 – 0.13 | -0.13 – 0.32 | -0.12 | 0.86 | 0.905 | 0.392 |
| Time in days | -0.00 | 0.00 | -0.05 | 0.05 | -0.01 – 0.00 | -0.15 – 0.05 | -0.94 | -0.94 | 0.348 | 0.348 |
| *APOE4* Group [carrier] | 0.00 | 0.01 | -0.05 | 0.12 | -0.02 – 0.02 | -0.29 – 0.19 | 0.01 | -0.39 | 0.994 | 0.694 |
| Age in years | 0.00 | 0.00 | 0.06 | 0.06 | -0.00 – 0.00 | -0.06 – 0.18 | 0.97 | 0.97 | 0.332 | 0.332 |
| Sex [female] | -0.01 | 0.01 | -0.14 | 0.13 | -0.03 – 0.01 | -0.38 – 0.11 | -1.07 | -1.07 | 0.284 | 0.284 |
| Education in years | 0.00 | 0.00 | 0.04 | 0.06 | -0.00 – 0.00 | -0.07 – 0.16 | 0.72 | 0.72 | 0.471 | 0.471 |
| Time in days × *APOE4* Group [carrier] | 0.01 | 0.01 | 0.09 | 0.09 | -0.01 – 0.02 | -0.09 – 0.27 | 1.02 | 1.02 | 0.311 | 0.311 |
| **Random Effects** | | | | | | | | | | |
| $\sigma^2$ | 0.00 | | | | | | | | | |
| $\tau_{00\ subj}$ | 0.00 | | | | | | | | | |
| ICC | 0.25 | | | | | | | | | |
| $N_{subj}$ | 152 | | | | | | | | | |
| Observations | 457 | | | | | | | | | |
| Marginal $R^2$ / Conditional $R^2$ | 0.014 / 0.265 | | | | | | | | | |

FC = functional connectivity. MTL = medial temporal lobe. PMC = posteromedial cortex. CI = 95% confidence interval.

#### Supplementary Tab. 7. Linear model of effects of covariates on retrieval-FC within MTL over time

| | Functional connectivity strength during retrieval within MTL | | | | | | | | | |
|---|---|---|---|---|---|---|---|---|---|---|
| *Predictors* | *Estimates* | *std. Error* | *std. Beta* | *standardized std. Error* | *CI* | *standardized CI* | *Statistic* | *std. Statistic* | *p* | *std. p* |
| (Intercept) | 0.46 | 0.08 | 0.29 | 0.13 | 0.31 – 0.61 | 0.04 – 0.54 | 6.01 | 2.26 | **<0.001** | **0.025** |
| Time in days | 0.00 | 0.00 | 0.01 | 0.04 | -0.01 – 0.01 | -0.07 – 0.09 | 0.28 | 0.28 | 0.779 | 0.779 |
| *APOE4* Group [carrier] | 0.00 | 0.01 | 0.05 | 0.14 | -0.02 – 0.03 | -0.22 – 0.32 | 0.44 | 0.36 | 0.658 | 0.722 |
| Age in years | -0.00 | 0.00 | -0.18 | 0.07 | -0.01 – -0.00 | -0.32 – -0.05 | -2.73 | -2.73 | **0.007** | **0.007** |
| Sex [female] | -0.03 | 0.01 | -0.47 | 0.14 | -0.06 – -0.01 | -0.75 – -0.19 | -3.35 | -3.35 | **0.001** | **0.001** |
| Education in years | 0.00 | 0.00 | 0.02 | 0.07 | -0.00 – 0.00 | -0.11 – 0.15 | 0.35 | 0.35 | 0.730 | 0.730 |
| Time in days × *APOE4* Group [carrier] | 0.00 | 0.01 | 0.03 | 0.07 | -0.01 – 0.01 | -0.11 – 0.16 | 0.41 | 0.41 | 0.680 | 0.680 |
| **Random Effects** | | | | | | | | | | |
| $\sigma^2$ | 0.00 | | | | | | | | | |
| $\tau_{00\ subj}$ | 0.00 | | | | | | | | | |
| ICC | 0.55 | | | | | | | | | |
| $N_{subj}$ | 152 | | | | | | | | | |
| Observations | 457 | | | | | | | | | |
| Marginal $R^2$ / Conditional $R^2$ | 0.076 / 0.581 | | | | | | | | | |

FC = functional connectivity. MTL = medial temporal lobe. CI = 95% confidence interval.

#### Supplementary Tab. 8. Linear model of effects of covariates on retrieval-FC within PMC over time

| | Functional connectivity strength during retrieval within PMC | | | | | | | | | |
|---|---|---|---|---|---|---|---|---|---|---|
| *Predictors* | *Estimates* | *std. Error* | *std. Beta* | *standardized std. Error* | *CI* | *standardized CI* | *Statistic* | *std. Statistic* | *p* | *std. p* |
| (Intercept) | 0.41 | 0.08 | 0.48 | 0.12 | 0.25 – 0.57 | 0.24 – 0.72 | 5.02 | 3.92 | **<0.001** | **<0.001** |
| Time in days | 0.00 | 0.00 | 0.04 | 0.04 | -0.00 – 0.01 | -0.04 – 0.12 | 0.89 | 0.89 | 0.375 | 0.375 |

| | | | | | | | | | | |
|---|---|---|---|---|---|---|---|---|---|---|
| *APOE4* Group [carrier] | -0.01 | 0.01 | -0.01 | 0.13 | -0.03 – 0.02 | -0.26 – 0.25 | -0.47 | -0.05 | 0.640 | 0.960 |
| Age in years | 0.00 | 0.00 | 0.08 | 0.06 | -0.00 – 0.00 | -0.05 – 0.20 | 1.19 | 1.19 | 0.237 | 0.237 |
| Sex [female] | -0.06 | 0.01 | -0.68 | 0.13 | -0.08 – -0.03 | -0.95 – -0.42 | -5.10 | -5.10 | **<0.001** | **<0.001** |
| Education in years | -0.00 | 0.00 | -0.05 | 0.06 | -0.00 – 0.00 | -0.17 – 0.08 | -0.78 | -0.78 | 0.437 | 0.437 |
| Time in days × *APOE4* Group [carrier] | -0.01 | 0.01 | -0.11 | 0.07 | -0.02 – 0.00 | -0.25 – 0.03 | -1.49 | -1.49 | 0.137 | 0.137 |
| **Random Effects** | | | | | | | | | | |
| $\sigma^2$ | 0.00 | | | | | | | | | |
| $\tau_{00\ subj}$ | 0.00 | | | | | | | | | |
| ICC | 0.47 | | | | | | | | | |
| $N_{subj}$ | 152 | | | | | | | | | |
| Observations | 457 | | | | | | | | | |
| Marginal $R^2$ / Conditional $R^2$ | 0.122 / 0.537 | | | | | | | | | |

FC = functional connectivity. PMC = posteromedial cortex. CI = 95% confidence interval.

#### Supplementary Tab. 9. Linear model of effects of covariates on retrieval-FC between MTL and PMC over time

| | Functional connectivity strength during retrieval between MTL and PMC | | | | | | | | | |
|---|---|---|---|---|---|---|---|---|---|---|
| *Predictors* | *Estimates* | *std. Error* | *std. Beta* | *standardized std. Error* | *CI* | *standardized CI* | *Statistic* | *std. Statistic* | *p* | *std. p* |
| (Intercept) | 0.15 | 0.05 | 0.17 | 0.12 | 0.05 – 0.25 | -0.06 – 0.40 | 2.88 | 1.44 | **0.005** | 0.152 |
| Time in days | -0.00 | 0.00 | -0.00 | 0.05 | -0.01 – 0.01 | -0.10 – 0.09 | -0.06 | -0.06 | 0.952 | 0.952 |
| *APOE4* Group [carrier] | -0.00 | 0.01 | -0.03 | 0.12 | -0.02 – 0.01 | -0.28 – 0.21 | -0.37 | -0.26 | 0.712 | 0.793 |
| Age in years | -0.00 | 0.00 | -0.10 | 0.06 | -0.00 – 0.00 | -0.22 – 0.02 | -1.59 | -1.59 | 0.113 | 0.113 |
| Sex [female] | -0.01 | 0.01 | -0.24 | 0.13 | -0.03 – 0.00 | -0.49 – 0.02 | -1.85 | -1.85 | 0.067 | 0.067 |
| Education in years | 0.00 | 0.00 | 0.04 | 0.06 | -0.00 – 0.00 | -0.08 – 0.16 | 0.68 | 0.68 | 0.499 | 0.499 |
| Time in days × *APOE4* Group [carrier] | -0.00 | 0.01 | -0.03 | 0.09 | -0.01 – 0.01 | -0.21 – 0.14 | -0.38 | -0.38 | 0.706 | 0.706 |
| **Random Effects** | | | | | | | | | | |

| | |
|---|---|
| $\sigma^2$ | 0.00 |
| $\tau_{00\ subj}$ | 0.00 |
| ICC | 0.29 |
| $N_{subj}$ | 152 |
| Observations | 457 |
| Marginal $R^2$ / Conditional $R^2$ | 0.021 / 0.306 |

FC = functional connectivity. MTL = medial temporal lobe. PMC = posteromedial cortex. CI = 95% confidence interval.

#### Supplementary Tab. 10. Linear model of effects of rsFC and covariates on amyloid-PET burden

| | **Global neocortical amyloid-PET burden** | | | | | | | |
|---|---|---|---|---|---|---|---|---|
| *Predictors* | *Estimates* | *std. Error* | *std. Beta* | *standardized std. Error* | *CI* | *standardized CI* | *Statistic* | *p* |
| (Intercept) | -0.06 | 0.12 | -0.44 | 0.18 | -0.30 – 0.18 | -0.79 – -0.10 | -0.50 | 0.621 |
| Slope of rsFC within MTL | -6.92 | 4.25 | -0.14 | 0.09 | -15.33 – 1.48 | -0.31 – 0.03 | -1.63 | 0.106 |
| *APOE4* Group [carrier] | 0.07 | 0.02 | 0.69 | 0.17 | 0.03 – 0.10 | 0.36 – 1.03 | 4.07 | **<0.001** |
| Slope of rsFC within PMC | 3.81 | 9.89 | 0.04 | 0.11 | -15.74 – 23.36 | -0.18 – 0.26 | 0.39 | 0.701 |
| Slope of rsFC between MTL and PMC | -3.98 | 3.84 | -0.11 | 0.11 | -11.56 – 3.61 | -0.33 – 0.10 | -1.04 | 0.302 |
| Age in years | 0.00 | 0.00 | 0.15 | 0.08 | -0.00 – 0.00 | 0.00 – 0.32 | 1.86 | 0.065 |
| Sex [female] | 0.02 | 0.02 | 0.21 | 0.20 | -0.02 – 0.06 | -0.159– 0.61 | 1.04 | 0.302 |
| Education in years | -0.00 | 0.00 | -0.09 | 0.07 | -0.01 – 0.00 | -0.22 – 0.04 | -1.33 | 0.184 |
| Time baseline to PET | 0.00 | 0.00 | 0.09 | 0.08 | -0.00 – 0.00 | -0.06 – 0.24 | 1.14 | 0.257 |
| Slope of rsFC within MTL × *APOE4* Group [carrier] | -6.11 | 9.67 | -0.12 | 0.20 | -25.22 – 13.00 | -0.52 – 0.27 | -0.63 | 0.529 |
| Slope of rsFC within PMC × *APOE4* Group [carrier] | -36.95 | 17.60 | -0.42 | 0.20 | -71.74 – -2.16 | -0.81 – -0.02 | -2.10 | **0.038** |
| Slope of rsFC between MTL and PMC × *APOE4* Group [carrier] | 11.72 | 7.63 | 0.34 | 0.22 | -3.36 – 26.80 | -0.10 – 0.77 | 1.54 | 0.127 |
| Observations | 152 | | | | | | | |

| | |
|---|---|
| $R^2$ / $R^2$ adjusted | 0.235 / 0.175 |

rsFC = resting-state functional connectivity. MTL = medial temporal lobe. PMC = posteromedial cortex. CI = 95% confidence interval.

#### Supplementary Tab. 11. Linear model of effects of rsFC and covariates on amyloid-PET burden in *APOE4* carriers

| | Global neocortical amyloid-PET burden | | | | | | | |
|---|---|---|---|---|---|---|---|---|
| *Predictors* | *Estimates* | *std. Error* | *std. Beta* | *standardized std. Error* | *CI* | *standardized CI* | *Statistic* | *p* |
| (Intercept) | -0.13 | 0.25 | -0.06 | 0.23 | -0.63 – 0.37 | -0.52 – 0.40 | -0.53 | 0.602 |
| Slope of rsFC within MTL | -13.34 | 8.55 | -0.23 | 0.15 | -30.49 – 3.82 | -0.52 – 0.07 | -1.56 | 0.125 |
| Slope of rsFC within PMC | -31.51 | 15.99 | -0.35 | 0.18 | -63.61 – 0.60 | -0.70 – 0.01 | -1.97 | 0.054 |
| Slope of rsFC between MTL and PMC | 7.94 | 6.73 | 0.20 | 0.17 | -5.58 – 21.46 | -0.14 – 0.54 | 1.18 | 0.244 |
| Age in years | 0.01 | 0.00 | 0.21 | 0.14 | -0.00 – 0.01 | -0.06 – 0.48 | 1.53 | 0.131 |
| Sex [female] | 0.01 | 0.03 | 0.10 | 0.30 | -0.05 – 0.08 | -0.50 – 0.69 | 0.32 | 0.747 |
| Education in years | -0.00 | 0.00 | -0.13 | 0.13 | -0.01 – 0.00 | -0.39 – 0.13 | -0.99 | 0.326 |
| Time baseline to PET | 0.00 | 0.00 | 0.18 | 0.13 | -0.00 – 0.00 | -0.09 – 0.45 | 1.35 | 0.183 |
| Observations | 59 | | | | | | | |
| $R^2$ / $R^2$ adjusted | 0.169 / 0.054 | | | | | | | |

rsFC = resting-state functional connectivity. MTL = medial temporal lobe. PMC = posteromedial cortex. CI = 95% confidence interval.

#### Supplementary Tab. 12. Linear model of effects of rsFC and covariates on amyloid-PET burden in APOE4 non-carriers

| | Global neocortical amyloid-PET burden | | | | | | | |
|---|---|---|---|---|---|---|---|---|
| *Predictors* | *Estimates* | *std. Error* | *std. Beta* | *standardized std. Error* | *CI* | *standardized CI* | *Statistic* | *p* |
| (Intercept) | -0.05 | 0.13 | -0.27 | 0.21 | -0.30 – 0.21 | -0.68 – 0.15 | -0.36 | 0.718 |
| Slope of rsFC within MTL | -6.02 | 4.82 | -0.16 | 0.13 | -15.61 – 3.57 | -0.42 – 0.10 | -1.25 | 0.215 |
| Slope of rsFC within PMC | 0.75 | 10.54 | 0.01 | 0.14 | -20.21 – 21.70 | -0.27 – 0.29 | 0.07 | 0.944 |

| | | | | | | | | |
|---|---|---|---|---|---|---|---|---|
| Slope of rsFC between MTL and PMC | -3.31 | 4.09 | -0.12 | 0.15 | -11.44 – 4.81 | -0.41 – 0.17 | -0.81 | 0.420 |
| Age in years | 0.00 | 0.00 | 0.18 | 0.11 | -0.00 – 0.01 | -0.03 – 0.39 | 1.66 | 0.100 |
| Sex [female] | 0.03 | 0.02 | 0.39 | 0.26 | -0.01 – 0.07 | -0.14 – 0.91 | 1.47 | 0.146 |
| Education in years | -0.00 | 0.00 | -0.06 | 0.11 | -0.01 – 0.00 | -0.29 – 0.16 | -0.55 | 0.586 |
| Time baseline to PET | 0.00 | 0.00 | 0.03 | 0.11 | -0.00 – 0.00 | -0.18 – 0.24 | 0.32 | 0.750 |
| Observations | 93 | | | | | | | |
| $R^2$ / $R^2$ adjusted | 0.098 / 0.023 | | | | | | | |

rsFC = resting-state functional connectivity. MTL = medial temporal lobe. PMC = posteromedial cortex. CI = 95% confidence interval.

#### Supplementary Tab. 13. Linear model of effects of encoding-FC and covariates on amyloid-PET burden

| | Global neocortical amyloid-PET burden | | | | | | | |
|---|---|---|---|---|---|---|---|---|
| *Predictors* | *Estimates* | *std. Error* | *std. Beta* | *standardized std. Error* | *CI* | *standardized CI* | *Statistic* | *p* |
| (Intercept) | -0.09 | 0.12 | -0.52 | 0.16 | -0.33 – 0.16 | -0.83 – -0.21 | -0.71 | 0.480 |
| Slope of encoding-FC within MTL | 0.12 | 0.68 | 0.02 | 0.10 | -1.22 – 1.46 | -0.18 – 0.21 | 0.17 | 0.863 |
| *APOE4* Group [carrier] | 0.08 | 0.02 | 0.78 | 0.16 | 0.04 – 0.11 | 0.46 – 1.09 | 4.90 | **<0.001** |
| Slope of encoding-FC within PMC | -1.64 | 4.17 | -0.04 | 0.11 | -9.89 – 6.61 | -0.26 – 0.17 | -0.39 | 0.695 |
| Slope of encoding-FC between MTL and PMC | -1.66 | 3.26 | -0.05 | 0.10 | -8.10 – 4.78 | -0.25 – 0.15 | -0.51 | 0.612 |
| Age in years | 0.00 | 0.00 | 0.17 | 0.08 | 0.00 – 0.01 | 0.01 – 0.33 | 2.07 | **0.040** |
| Sex [female] | 0.03 | 0.02 | 0.33 | 0.17 | -0.00 – 0.07 | -0.01 – 0.68 | 1.90 | 0.060 |
| Education in years | -0.00 | 0.00 | -0.08 | 0.08 | -0.01 – 0.00 | -0.24 – 0.07 | -1.08 | 0.284 |
| Time baseline to PET | 0.00 | 0.00 | 0.09 | 0.08 | -0.00 – 0.00 | -0.07 – 0.24 | 1.12 | 0.263 |
| Slope of encoding-FC within MTL × *APOE4* Group [carrier] | 0.09 | 1.17 | 0.01 | 0.17 | -2.22 – 2.40 | -0.32 – 0.34 | 0.08 | 0.938 |
| Slope of encoding-FC within PMC × *APOE4* Group [carrier] | -2.21 | 7.12 | -0.06 | 0.18 | -16.29 – 11.88 | -0.42 – 0.31 | -0.31 | 0.757 |

| | | | | | | | | |
|---|---|---|---|---|---|---|---|---|
| Slope of encoding-FC between MTL and PMC × *APOE4* Group [carrier] | 3.99 | 6.25 | 0.13 | 0.20 | -8.38 – 16.35 | -0.26 – 0.51 | 0.64 | 0.525 |
| Observations | 152 | | | | | | | |
| $R^2$ / $R^2$ adjusted | 0.189 / 0.125 | | | | | | | |

FC = functional connectivity. MTL = medial temporal lobe. PMC = posteromedial cortex. CI = 95% confidence interval.

#### Supplementary Tab. 14. Linear model of effects of retrieval-FC and covariates on amyloid-PET burden

| | Global neocortical amyloid-PET burden | | | | | | | |
|---|---|---|---|---|---|---|---|---|
| *Predictors* | *Estimates* | *std. Error* | *std. Beta* | *standardized std. Error* | *CI* | *standardized CI* | *Statistic* | *p* |
| (Intercept) | -0.10 | 0.12 | -0.49 | 0.15 | -0.34 – 0.14 | -0.80 – -0.19 | -0.81 | 0.421 |
| Slope of retrieval-FC within MTL | -9.13 | 10.39 | -0.09 | 0.10 | -29.68 – 11.42 | -0.28 – 0.11 | -0.88 | 0.381 |
| *APOE4* Group [carrier] | 0.07 | 0.02 | 0.76 | 0.16 | 0.04 – 0.10 | 0.45 – 1.08 | 4.80 | **<0.001** |
| Slope of retrieval-FC within PMC | -1.42 | 1.18 | -0.12 | 0.10 | -3.75 – 0.91 | -0.31 – 0.08 | -1.21 | 0.230 |
| Slope of retrieval-FC between MTL and PMC | 3.69 | 5.35 | 0.08 | 0.11 | -6.89 – 14.27 | -0.14 – 0.29 | 0.69 | 0.492 |
| Age in years | 0.00 | 0.00 | 0.17 | 0.08 | 0.00 – 0.01 | 0.01 – 0.33 | 2.13 | **0.035** |
| Sex [female] | 0.03 | 0.02 | 0.29 | 0.17 | -0.01 – 0.06 | -0.05 – 0.63 | 1.67 | 0.097 |
| Education in years | -0.00 | 0.00 | -0.07 | 0.08 | -0.01 – 0.00 | -0.23 – 0.09 | -0.89 | 0.373 |
| Time baseline to PET | 0.00 | 0.00 | 0.09 | 0.08 | -0.00 – 0.00 | -0.06 – 0.25 | 1.16 | 0.249 |
| Slope of retrieval-FC within MTL × *APOE4* Group [carrier] | 22.16 | 17.70 | 0.21 | 0.17 | -12.83 – 57.16 | -0.12 – 0.54 | 1.25 | 0.213 |
| Slope of retrieval-FC within PMC × *APOE4* Group [carrier] | 1.28 | 2.14 | 0.11 | 0.18 | -2.95 – 5.51 | -0.25 – 0.46 | 0.60 | 0.550 |
| Slope of retrieval-FC between MTL and PMC × *APOE4* Group [carrier] | -0.09 | 8.17 | -0.00 | 0.17 | -16.25 – 16.07 | -0.34 – 0.33 | -0.01 | 0.991 |
| Observations | 152 | | | | | | | |
| $R^2$ / $R^2$ adjusted | 0.196 / 0.133 | | | | | | | |

FC = functional connectivity. MTL = medial temporal lobe. PMC = posteromedial cortex. CI = 95% confidence interval.

#### Supplementary Tab. 15. Linear model of effects of encoding-FC and covariates on tau-PET burden

| | Entorhinal tau-PET burden | | | | | | | |
|---|---|---|---|---|---|---|---|---|
| *Predictors* | *Estimates* | *std. Error* | *std. Beta* | *standardized std. Error* | *CI* | *standardized CI* | *Statistic* | *p* |
| (Intercept) | -0.30 | 0.12 | -0.53 | 0.16 | -0.53 – -0.08 | -0.85 – -0.22 | -2.63 | **0.009** |
| Slope of encoding-FC within MTL | -0.22 | 0.63 | -0.04 | 0.10 | -1.48 – 1.03 | -0.23 – 0.16 | -0.36 | 0.723 |
| *APOE4* Group [carrier] | 0.03 | 0.01 | 0.30 | 0.16 | -0.00 – 0.05 | -0.03 – 0.62 | 1.82 | 0.071 |
| Slope of encoding-FC within PMC | 2.78 | 3.90 | 0.08 | 0.11 | -4.93 – 10.49 | -0.14 – 0.30 | 0.71 | 0.478 |
| Slope of encoding-FC between MTL and PMC | -1.46 | 3.04 | -0.05 | 0.11 | -7.48 – 4.56 | -0.26 – 0.16 | -0.48 | 0.633 |
| Age in years | 0.00 | 0.00 | 0.20 | 0.08 | 0.00 – 0.01 | 0.04 – 0.37 | 2.44 | **0.016** |
| Sex [female] | 0.05 | 0.02 | 0.61 | 0.18 | 0.02 – 0.08 | 0.25 – 0.97 | 3.39 | **0.001** |
| Education in years | 0.00 | 0.00 | 0.09 | 0.08 | -0.00 – 0.01 | -0.07 – 0.25 | 1.12 | 0.265 |
| Time baseline to PET | 0.00 | 0.00 | 0.02 | 0.08 | -0.00 – 0.00 | -0.14 – 0.18 | 0.25 | 0.806 |
| Slope of encoding-FC within MTL × *APOE4* Group [carrier] | 2.16 | 1.09 | 0.34 | 0.17 | 0.00 – 4.32 | 0.00 – 0.68 | 1.98 | **0.0497** |
| Slope of encoding-FC within PMC × *APOE4* Group [carrier] | -4.04 | 6.66 | -0.12 | 0.19 | -17.21 – 9.12 | -0.49 – 0.26 | -0.61 | 0.545 |
| Slope of encoding-FC between MTL and PMC × *APOE4* Group [carrier] | 4.36 | 5.85 | 0.15 | 0.20 | -7.20 – 15.92 | -0.25 – 0.55 | 0.75 | 0.457 |
| Observations | 152 | | | | | | | |
| $R^2$ / $R^2$ adjusted | 0.141 / 0.074 | | | | | | | |

FC = functional connectivity. MTL = medial temporal lobe. PMC = posteromedial cortex. CI = 95% confidence interval.

#### Supplementary Tab. 16. Linear model of effects of encoding-FC and covariates on tau-PET burden in *APOE4* carriers

| | Entorhinal tau-PET burden | | | | | | | |
|---|---|---|---|---|---|---|---|---|
| *Predictors* | *Estimates* | *std. Error* | *std. Beta* | *standardized std. Error* | *CI* | *standardized CI* | *Statistic* | *p* |

| | | | | | | | | |
|---|---|---|---|---|---|---|---|---|
| (Intercept) | -0.38 | 0.22 | -0.37 | 0.23 | -0.83 – 0.07 | -0.82 – 0.09 | -1.71 | 0.093 |
| Slope of encoding-FC within MTL | 1.94 | 1.04 | 0.27 | 0.14 | -0.14 – 4.02 | -0.02 – 0.55 | 1.88 | 0.066 |
| Slope of encoding-FC within PMC | -0.38 | 6.25 | -0.01 | 0.17 | -12.92 – 12.15 | -0.36 – 0.33 | -0.06 | 0.951 |
| Slope of encoding-FC between MTL and PMC | 3.54 | 5.63 | 0.11 | 0.18 | -7.75 – 14.84 | -0.24 – 0.46 | 0.63 | 0.532 |
| Age in years | 0.01 | 0.00 | 0.25 | 0.14 | -0.00 – 0.01 | -0.04 – 0.53 | 1.75 | 0.087 |
| Sex [female] | 0.05 | 0.03 | 0.57 | 0.29 | -0.00 – 0.11 | -0.01 – 1.15 | 1.97 | 0.054 |
| Education in years | 0.00 | 0.00 | 0.04 | 0.13 | -0.01 – 0.01 | -0.22 – 0.31 | 0.33 | 0.739 |
| Time baseline to PET | 0.00 | 0.00 | 0.06 | 0.14 | -0.00 – 0.00 | -0.22 – 0.34 | 0.44 | 0.663 |
| Observations | 59 | | | | | | | |
| $R^2$ / $R^2$ adjusted | 0.152 / 0.036 | | | | | | | |

FC = functional connectivity. MTL = medial temporal lobe. PMC = posteromedial cortex. CI = 95% confidence interval.

#### Supplementary Tab. 17. Linear model of effects of encoding-FC and covariates on tau-PET burden in *APOE4* non-carriers

| | Entorhinal tau-PET burden | | | | | | | |
|---|---|---|---|---|---|---|---|---|
| *Predictors* | *Estimates* | *std. Error* | *std. Beta* | *standardized std. Error* | *CI* | *standardized CI* | *Statistic* | *p* |
| (Intercept) | -0.27 | 0.14 | -0.44 | 0.19 | -0.54 – 0.00 | -0.82 – -0.06 | -1.95 | 0.054 |
| Slope of encoding-FC within MTL | -0.24 | 0.61 | -0.04 | 0.11 | -1.46 – 0.98 | -0.25 – 0.17 | -0.40 | 0.693 |
| Slope of encoding-FC within PMC | 2.65 | 3.76 | 0.08 | 0.11 | -4.83 – 10.13 | -0.14 – 0.30 | 0.70 | 0.483 |
| Slope of encoding-FC between MTL and PMC | -1.37 | 2.91 | -0.05 | 0.11 | -7.16 – 4.42 | -0.27 – 0.17 | -0.47 | 0.639 |
| Age in years | 0.00 | 0.00 | 0.19 | 0.11 | -0.00 – 0.01 | -0.02 – 0.40 | 1.77 | 0.081 |
| Sex [female] | 0.05 | 0.02 | 0.64 | 0.24 | 0.01 – 0.09 | 0.17 – 1.11 | 2.70 | **0.008** |
| Education in years | 0.00 | 0.00 | 0.11 | 0.10 | -0.00 – 0.01 | -0.09 – 0.32 | 1.09 | 0.281 |
| Time baseline to PET | 0.00 | 0.00 | 0.01 | 0.10 | -0.00 – 0.00 | -0.20 – 0.22 | 0.07 | 0.942 |
| Observations | 93 | | | | | | | |

| | |
|---|---|
| $R^2$ / $R^2$ adjusted | 0.117 / 0.045 |

FC = functional connectivity. MTL = medial temporal lobe. PMC = posteromedial cortex. CI = 95% confidence interval.

#### Supplementary Tab. 18. Linear model of effects of retrieval-FC and covariates on tau-PET burden

| | Entorhinal tau-PET burden | | | | | | | |
|---|---|---|---|---|---|---|---|---|
| *Predictors* | *Estimates* | *std. Error* | *std. Beta* | *standardized std. Error* | *CI* | *standardized CI* | *Statistic* | *p* |
| (Intercept) | -0.32 | 0.11 | -0.52 | 0.15 | -0.54 – -0.10 | -0.82 – -0.21 | -2.87 | **0.005** |
| Slope of retrieval-FC within MTL | -6.44 | 9.41 | -0.07 | 0.10 | -25.05 – 12.17 | -0.26 – 0.13 | -0.68 | 0.495 |
| *APOE4* Group [carrier] | 0.03 | 0.01 | 0.32 | 0.16 | 0.00 – 0.06 | 0.00 – 0.63 | 1.99 | **0.048** |
| Slope of retrieval-FC within PMC | -3.78 | 1.07 | -0.35 | 0.10 | -5.89 – -1.67 | -0.54 – -0.15 | -3.54 | **0.001** |
| Slope of retrieval-FC between MTL and PMC | 12.31 | 4.85 | 0.28 | 0.11 | 2.73 – 21.89 | 0.06 – 0.50 | 2.54 | **0.012** |
| Age in years | 0.00 | 0.00 | 0.22 | 0.08 | 0.00 – 0.01 | 0.06 – 0.38 | 2.66 | **0.009** |
| Sex [female] | 0.05 | 0.02 | 0.61 | 0.17 | 0.02 – 0.08 | 0.27 – 0.96 | 3.54 | **0.001** |
| Education in years | 0.00 | 0.00 | 0.09 | 0.08 | -0.00 – 0.01 | -0.07 – 0.24 | 1.15 | 0.253 |
| Time baseline to PET | 0.00 | 0.00 | 0.04 | 0.08 | -0.00 – 0.00 | -0.12 – 0.19 | 0.48 | 0.635 |
| Slope of retrieval-FC within MTL × *APOE4* Group [carrier] | 29.70 | 16.03 | 0.31 | 0.17 | -2.00 – 61.39 | -0.02 – 0.63 | 1.85 | 0.066 |
| Slope of retrieval-FC within PMC × *APOE4* Group [carrier] | 6.32 | 1.94 | 0.58 | 0.18 | 2.49 – 10.15 | 0.23 – 0.94 | 3.26 | **0.001** |
| Slope of retrieval-FC between MTL and PMC × *APOE4* Group [carrier] | -14.03 | 7.40 | -0.32 | 0.17 | -28.66 – 0.60 | -0.65 – 0.01 | -1.90 | 0.060 |
| Observations | 152 | | | | | | | |
| $R^2$ / $R^2$ adjusted | 0.201 / 0.139 | | | | | | | |

FC = functional connectivity. MTL = medial temporal lobe. PMC = posteromedial cortex. CI = 95% confidence interval.

#### Supplementary Tab. 19. Linear model of effects of retrieval-FC and covariates on tau-PET burden in *APOE4* carriers

| Predictors | Entorhinal tau-PET burden | | | | | | | |
|---|---|---|---|---|---|---|---|---|
| | Estimates | std. Error | std. Beta | standardized std. Error | CI | standardized CI | Statistic | p |
| (Intercept) | -0.39 | 0.22 | -0.27 | 0.23 | -0.84 – 0.06 | -0.73 – 0.19 | -1.73 | 0.089 |
| Slope of retrieval-FC within MTL | 22.31 | 15.99 | 0.20 | 0.14 | -9.80 – 54.41 | -0.09 – 0.48 | 1.39 | 0.169 |
| Slope of retrieval-FC within PMC | 2.77 | 1.90 | 0.20 | 0.14 | -1.04 – 6.58 | -0.07 – 0.47 | 1.46 | 0.151 |
| Slope of retrieval-FC between MTL and PMC | -1.10 | 6.58 | -0.02 | 0.14 | -14.32 – 12.12 | -0.30 – 0.25 | -0.17 | 0.867 |
| Age in years | 0.01 | 0.00 | 0.26 | 0.14 | -0.00 – 0.01 | -0.02 – 0.54 | 1.83 | 0.073 |
| Sex [female] | 0.04 | 0.03 | 0.42 | 0.29 | -0.02 – 0.09 | -0.17 – 1.00 | 1.42 | 0.161 |
| Education in years | 0.00 | 0.00 | 0.05 | 0.13 | -0.01 – 0.01 | -0.22 – 0.32 | 0.38 | 0.704 |
| Time baseline to PET | 0.00 | 0.00 | 0.05 | 0.14 | -0.00 – 0.00 | -0.22 – 0.33 | 0.39 | 0.699 |
| Observations | 59 | | | | | | | |
| $R^2$ / $R^2$ adjusted | 0.128 / 0.008 | | | | | | | |

FC = functional connectivity. MTL = medial temporal lobe. PMC = posteromedial cortex. CI = 95% confidence interval.

#### Supplementary Tab. 20. Linear model of effects of retrieval-FC and covariates on tau-PET burden in *APOE4* non-carriers

| Predictors | Entorhinal tau-PET burden | | | | | | | |
|---|---|---|---|---|---|---|---|---|
| | Estimates | std. Error | std. Beta | standardized std. Error | CI | standardized CI | Statistic | p |
| (Intercept) | -0.30 | 0.13 | -0.53 | 0.18 | -0.55 – -0.05 | -0.88 – -0.18 | -2.35 | **0.021** |
| Slope of retrieval-FC within MTL | -6.02 | 8.73 | -0.07 | 0.10 | -23.38 – 11.34 | -0.27 – 0.13 | -0.69 | 0.493 |
| Slope of retrieval-FC within PMC | -3.84 | 0.97 | -0.41 | 0.10 | -5.77 – -1.90 | -0.62 – -0.20 | -3.94 | **<0.001** |
| Slope of retrieval-FC between MTL and PMC | 11.97 | 4.46 | 0.29 | 0.11 | 3.10 – 20.84 | 0.08 – 0.51 | 2.68 | **0.009** |
| Age in years | 0.00 | 0.00 | 0.21 | 0.10 | 0.00 – 0.01 | 0.01 – 0.41 | 2.04 | **0.044** |
| Sex [female] | 0.06 | 0.02 | 0.77 | 0.22 | 0.03 – 0.10 | 0.33 – 1.20 | 3.50 | **0.001** |
| Education in years | 0.00 | 0.00 | 0.12 | 0.10 | -0.00 – 0.01 | -0.08 – 0.32 | 1.20 | 0.232 |

| | | | | | | | | |
|---|---|---|---|---|---|---|---|---|
| Time baseline to PET | 0.00 | 0.00 | 0.04 | 0.10 | -0.00 – 0.00 | -0.15 – 0.23 | 0.41 | 0.684 |
| Observations | 93 | | | | | | | |
| $R^2$ / $R^2$ adjusted | 0.257 / 0.196 | | | | | | | |

FC = functional connectivity. MTL = medial temporal lobe. PMC = posteromedial cortex. CI = 95% confidence interval.

#### Supplementary Tab. 21. Linear model of effects of covariates on the RBANS delayed memory index score over time

| | RBANS delayed memory index score | | | | | | | |
|---|---|---|---|---|---|---|---|---|
| *Predictors* | *Estimates* | *std. Error* | *std. Beta* | *standardized std. Error* | *CI* | *standardized CI* | *Statistic* | *p* |
| (Intercept) | 96.76 | 7.67 | -0.31 | 0.11 | 81.61 – 111.92 | -0.53 – -0.10 | 12.62 | **<0.001** |
| Time in days | 1.18 | 0.28 | 0.14 | 0.03 | 0.62 – 1.74 | 0.07 – 0.21 | 4.20 | **<0.001** |
| *APOE4* Group [carrier] | 0.88 | 1.00 | 0.10 | 0.11 | -1.09 – 2.85 | -0.13 – 0.33 | 0.88 | 0.380 |
| Age in years | -0.05 | 0.11 | -0.03 | 0.06 | -0.26 – 0.17 | -0.14 – 0.09 | -0.45 | 0.651 |
| Sex [female] | 3.54 | 1.04 | 0.41 | 0.12 | 1.47 – 5.60 | 0.17 – 0.65 | 3.39 | **0.001** |
| Education in years | 0.58 | 0.15 | 0.22 | 0.06 | 0.29 – 0.86 | 0.11 – 0.33 | 3.94 | **<0.001** |
| **Random Effects** | | | | | | | | |
| $\sigma^2$ | 42.59 | | | | | | | |
| $\tau_{00\ subj}$ | 25.23 | | | | | | | |
| $\tau 11_{\ subj.time}$ | 1.00 | | | | | | | |
| $\rho_{01\ subj}$ | 0.59 | | | | | | | |
| ICC | 0.37 | | | | | | | |
| $N_{\ subj}$ | 152 | | | | | | | |
| Observations | 575 | | | | | | | |
| Marginal $R^2$ / Conditional $R^2$ | 0.101 / 0.435 | | | | | | | |

RBANS = Repeatable Battery for the Assessment of Neuropsychological Status.

#### Supplementary Tab. 22. Linear model of effects of covariates on the fMRI-task corrected hit rate over time

| | fMRI retrieval task corrected hit rate |
|---|---|

| Predictors | Estimates | std. Error | std. Beta | standardized std. Error | CI | standardized CI | Statistic | p |
|---|---|---|---|---|---|---|---|---|
| (Intercept) | 0.70 | 0.20 | -0.13 | 0.12 | 0.31 – 1.10 | -0.37 – 0.12 | 3.50 | **0.001** |
| Time in days | -0.02 | 0.01 | -0.09 | 0.04 | -0.04 – -0.00 | -0.17 – -0.01 | -2.29 | **0.023** |
| *APOE4* Group [carrier] | -0.03 | 0.03 | -0.13 | 0.13 | -0.08 – 0.03 | -0.39 – 0.13 | -1.00 | 0.321 |
| Age in years | -0.00 | 0.00 | -0.02 | 0.06 | -0.01 – 0.00 | -0.14 – 0.11 | -0.23 | 0.816 |
| Sex [female] | 0.05 | 0.03 | 0.24 | 0.14 | -0.01 – 0.10 | -0.03 – 0.51 | 1.79 | 0.076 |
| Education in years | -0.00 | 0.00 | -0.01 | 0.06 | -0.01 – 0.01 | -0.14 – 0.12 | -0.15 | 0.879 |
| **Random Effects** | | | | | | | | |
| $\sigma^2$ | 0.02 | | | | | | | |
| $\tau_{00\ subj}$ | 0.01 | | | | | | | |
| ICC | 0.38 | | | | | | | |
| $N_{subj}$ | 152 | | | | | | | |
| Observations | 447 | | | | | | | |
| Marginal $R^2$ / Conditional $R^2$ | 0.027 / 0.393 | | | | | | | |

#### Supplementary Tab. 23. Linear model of effects of encoding-FC and covariates on slope of RBANS delayed memory index score

| | Slope of RBANS delayed memory index score | | | | | | | |
|---|---|---|---|---|---|---|---|---|
| Predictors | Estimates | std. Error | std. Beta | standardized std. Error | CI | standardized CI | Statistic | p |
| (Intercept) | -0.12 | 0.52 | -0.22 | 0.16 | -1.15 – 0.91 | -0.54 – 0.10 | -0.23 | 0.817 |
| Slope of encoding-FC within MTL | -2.87 | 2.90 | -0.10 | 0.10 | -8.61 – 2.87 | -0.30 – 0.10 | -0.99 | 0.325 |
| *APOE4* Group [carrier] | -0.00 | 0.07 | -0.00 | 0.17 | -0.13 – 0.13 | -0.33 – 0.33 | -0.01 | 0.991 |
| Slope of encoding-FC within PMC | -43.35 | 17.97 | -0.27 | 0.11 | -78.89 – -7.82 | -0.50 – -0.05 | -2.41 | **0.017** |
| Slope of encoding-FC between MTL and PMC | -18.84 | 14.02 | -0.14 | 0.11 | -46.55 – 8.87 | -0.36 – 0.07 | -1.34 | 0.181 |
| Age in years | -0.00 | 0.01 | -0.04 | 0.08 | -0.02 – 0.01 | -0.21 – 0.12 | -0.53 | 0.597 |
| Sex [female] | 0.12 | 0.07 | 0.31 | 0.18 | -0.02 – 0.27 | -0.05 – 0.68 | 1.72 | 0.088 |

| | | | | | | | | |
|---|---|---|---|---|---|---|---|---|
| Education in years | 0.02 | 0.01 | 0.15 | 0.08 | -0.00 – 0.04 | -0.01 – 0.32 | 1.86 | 0.064 |
| Time baseline to PET | 2.98 | 5.03 | 0.10 | 0.18 | -6.97 – 12.92 | -0.24 – 0.45 | 0.59 | 0.555 |
| Slope of encoding-FC within MTL × *APOE4* Group [carrier] | 46.41 | 30.68 | 0.29 | 0.19 | -14.23 – 107.06 | -0.09 – 0.68 | 1.51 | 0.133 |
| Slope of encoding-FC within PMC × *APOE4* Group [carrier] | 3.54 | 26.75 | 0.03 | 0.21 | -49.35 – 56.43 | -0.38 – 0.43 | 0.13 | 0.895 |
| Observations | 152 | | | | | | | |
| $R^2$ / $R^2$ adjusted | 0.093 / 0.029 | | | | | | | |

FC = functional connectivity. RBANS = Repeatable Battery for the Assessment of Neuropsychological Status. MTL = medial temporal lobe. PMC = posteromedial cortex. CI = 95% confidence interval.

#### Supplementary Tab. 24. Linear model of effects of encoding-FC and covariates on slope of episodic memory retrieval task corrected hit rate

| | Slope of fMRI retrieval task corrected hit rate | | | | | | | |
|---|---|---|---|---|---|---|---|---|
| *Predictors* | *Estimates* | *std. Error* | *std. Beta* | *standardized std. Error* | *CI* | *standardized CI* | *Statistic* | *p* |
| (Intercept) | 0.01 | 0.02 | -0.19 | 0.16 | -0.04 – 0.06 | -0.50 – 0.12 | 0.38 | 0.705 |
| Slope of encoding-FC within MTL | -0.06 | 0.13 | -0.05 | 0.10 | -0.33 – 0.20 | -0.24 – 0.15 | -0.47 | 0.640 |
| *APOE4* Group [carrier] | -0.00 | 0.00 | -0.18 | 0.16 | -0.01 – 0.00 | -0.50 – 0.14 | -1.10 | 0.273 |
| Slope of encoding-FC within PMC | -2.78 | 0.83 | -0.37 | 0.11 | -4.42 – -1.14 | -0.59 – -0.15 | -3.36 | **0.001** |
| Slope of encoding-FC between MTL and PMC | -1.63 | 0.65 | -0.26 | 0.10 | -2.91 – -0.35 | -0.47 – -0.06 | -2.52 | **0.013** |
| Age in years | -0.00 | 0.00 | -0.05 | 0.08 | -0.00 – 0.00 | -0.21 – 0.11 | -0.59 | 0.558 |
| Sex [female] | 0.01 | 0.00 | 0.39 | 0.18 | 0.00 – 0.01 | 0.04 – 0.74 | 2.17 | **0.031** |
| Education in years | 0.00 | 0.00 | 0.00 | 0.08 | -0.00 – 0.00 | -0.16 – 0.16 | 0.00 | 0.996 |
| Time baseline to PET | -0.08 | 0.23 | -0.06 | 0.17 | -0.54 – 0.38 | -0.40 – 0.28 | -0.34 | 0.733 |
| Slope of encoding-FC within MTL × *APOE4* Group [carrier] | 0.53 | 1.42 | 0.07 | 0.19 | -2.26 – 3.33 | -0.30 – 0.44 | 0.38 | 0.707 |

| | | | | | | | | |
|---|---|---|---|---|---|---|---|---|
| Slope of encoding-FC within PMC × *APOE4* Group [carrier] | -0.02 | 1.23 | -0.00 | 0.20 | -2.46 – 2.42 | -0.40 – 0.39 | -0.01 | 0.989 |
| Observations | 152 | | | | | | | |
| $R^2$ / $R^2$ adjusted | 0.148 / 0.087 | | | | | | | |

FC = functional connectivity. MTL = medial temporal lobe. PMC = posteromedial cortex. CI = 95% confidence interval.

#### Supplementary Tab. 25. Linear model of effects of rsFC and covariates on slope of RBANS delayed memory index score

| | Slope of RBANS delayed memory index score | | | | | | | |
|---|---|---|---|---|---|---|---|---|
| *Predictors* | *Estimates* | *std. Error* | *std. Beta* | *standardized std. Error* | *CI* | *standardized CI* | *Statistic* | *p* |
| (Intercept) | -0.15 | 0.51 | -0.14 | 0.17 | -1.16 – 0.87 | -0.48 – 0.20 | -0.28 | 0.777 |
| Slope of rsFC within MTL | 13.68 | 23.83 | 0.07 | 0.12 | -33.42 – 60.79 | -0.17 – 0.30 | 0.57 | 0.567 |
| *APOE4* Group [carrier] | -0.03 | 0.07 | -0.07 | 0.17 | -0.16 – 0.11 | -0.41 – 0.27 | -0.42 | 0.678 |
| Slope of rsFC within PMC | -100.58 | 52.88 | -0.28 | 0.15 | -205.13 – 3.97 | -0.57 – 0.01 | -1.90 | 0.059 |
| Slope of rsFC between MTL and PMC | 44.20 | 20.64 | 0.31 | 0.15 | 3.38 – 85.01 | 0.02 – 0.60 | 2.14 | **0.034** |
| Age in years | -0.00 | 0.01 | -0.04 | 0.08 | -0.02 – 0.01 | -0.21 – 0.12 | -0.50 | 0.619 |
| Sex [female] | 0.11 | 0.08 | 0.27 | 0.20 | -0.05 – 0.26 | -0.12 – 0.66 | 1.37 | 0.172 |
| Education in years | 0.02 | 0.01 | 0.17 | 0.08 | 0.00 – 0.04 | 0.01 – 0.34 | 2.05 | **0.043** |
| Time baseline to PET | -43.15 | 38.21 | -0.22 | 0.19 | -118.70 – 32.39 | -0.59 – 0.16 | -1.13 | 0.261 |
| Slope of rsFC within MTL × *APOE4* Group [carrier] | 104.58 | 78.08 | 0.29 | 0.22 | -49.78 – 258.93 | -0.14 – 0.71 | 1.34 | 0.183 |
| Slope of rsFC within PMC × *APOE4* Group [carrier] | -29.38 | 32.41 | -0.21 | 0.23 | -93.46 – 34.69 | -0.66 – 0.24 | -0.91 | 0.366 |
| Observations | 152 | | | | | | | |
| $R^2$ / $R^2$ adjusted | 0.089 / 0.025 | | | | | | | |

rsFC = resting-state functional connectivity. RBANS = Repeatable Battery for the Assessment of Neuropsychological Status. MTL = medial temporal lobe. PMC = posteromedial cortex. CI = 95% confidence interval.
